# A comparative assessment of self-limiting genetic control strategies for population suppression

**DOI:** 10.1101/2024.09.23.614516

**Authors:** Yue Han, Jackson Champer

## Abstract

Genetic control strategies are promising solutions for control of pest populations and invasive species. Methods utilizing repeated releases of males such as Sterile Insect Technique (SIT), Release of Insects carrying a Dominant Lethal (RIDL), self-limiting gene drives, and gene disruptors are highly controllable methods, ensuring biosafety. Although models of these strategies have been built, detailed comparisons are lacking, particularly for some of the newer strategies. Here, we conducted a thorough comparative assessment of self-limiting genetic control strategies by individual-based simulation models. Specifically, we find that repeated releases greatly enhance suppression power of weak and self-limiting gene drives, enabling population elimination with even low efficiency and high fitness costs. Moreover, dominant female sterility further strengthens self-limiting systems that can either use gene drive or disruptors that target genes without a mechanism to bias their own inheritance. Some of these strategies are highly persistent, resulting in relatively low release ratios even when released males suffer high fitness costs. To quantitively evaluate different strategies independent from ecological impact, we proposed constant-population genetic load, which achieves over 95% accuracy in predicting simulation outcomes for most strategies, though it is not as precise in a few frequency-dependent systems. Our results suggest that many new self-limiting strategies are safe, flexible, and more cost-effective than traditional SIT and RIDL, and thus have great potential for population suppression of insects and other pests.

## 1. Introduction

Insect pests such as fruit flies and mosquitoes post significant threats to agriculture and human health [1, 2]. Traditional pesticides can lead to environmental contamination [3], non-target species impact [4], and challenges such as rapidly developing resistance [5]. In recent years, alternative pest control strategies have shown great potential. Sterile insect technique (SIT) successfully reduces the populations of targeted species by releasing sterile males, which mate with females and produce no viable offspring [6–10]. Another strategy involves the use of *Wolbachia*, a symbiotic bacterium that causes cytoplasmic incompatibility [11], which has seen promising effects in controlling pest population and reducing diseases in multiple regions [12–15]. These methods avoid side effects of chemical pesticides, but also require substantial release sizes to suppress the population, resulting in considerable resource demands.

Genetic engineering approaches such as release of insects carrying a dominant lethal (RIDL) and female-specific RIDL (fsRIDL) have also been developed. These systems contain a dominant, conditional lethal gene element which can be repressed in the lab environment. When modified males are released into wild populations, this element will cause dominant lethality in their offspring (just female offspring for fsRIDL) [16]. RIDL has been developed in *Drosophila melanogaster* [17] and proved its feasibility in field tests of *Aedes aegypti* populations [18–20]. fsRIDL allows easy sex sorting and has been built in *Drosophila suzukii* [21], *Ceratitis capitata* [22, 23], *Aedes albopictus* [24], and seen success in *Ae. aegypti* field tests [25]. RIDL, fsRIDL, and genetic SIT strategies [26, 27] avoid harm caused by radiation or chemosterilization commonly seen in older SIT methods, increasing the chance of successful mating for released males. However, massive release sizes are still required to eliminate local populations.

Gene drive systems represent a more recent innovation in genetic pest control [28]. They are selfish genetic elements that bias mendelian inheritance to increase their frequency in the population, including suppression drives which reduce or eliminate the population, and modification drives which prevent transmission of diseases [29]. Various types of gene drives have been built in *D. melanogaster* [30–32], *D. suzukii* [33, 34], *Anopheles gambiae* [35–39], *Anopheles stephensi* [40–42], *Ae. aegypti* [43, 44] and *Plutella xylostella* [45]. Super-Mendelian inheritance was also achieved in mammals [46] and plants [47, 48].

Powerful self-sustaining suppression drives can spread through a population rapidly with even a small release size. Their frequency increases until reaching a stable equilibrium or completely suppressing the population. However, they also raise ecological and ethical concerns due to their uncontrolled spread [49–51]. Additionally, achieving high drive efficiency in some systems remains challenging [52–54]. Confined suppression drives are more controllable and require an introduction frequency above a certain threshold to increase their frequency and avoid elimination [55–60]. However, they can still persist in the environment if population elimination is not achieved. In contrast, self-limiting drives will themselves be eliminated over time [61–66]. These drives are more controllable and secure, but a single release often fails to provide adequate power for successful suppression.

Gene disruptors are genetic systems that target specific sites and disrupt gene functions. Unlike gene drives, a gene disruptor doesn’t increase its frequency in a population (with a few exceptions such as Y-linked X-shredders that can also be classified as gene drives). Instead, they rely on large or repeated releases to affect the population. These gene disruptors are naturally self-limiting and confined and thus less likely to provoke public opposition. Suppression gene disruptors with different targets have been modelled and built in *D. melanogaster* and *An. gambiae* [67–74]. Although gene disruptors never replicate themselves, they may act for multiple generations before being lost from the population. Therefore, gene disruptor strategies can be substantially more efficient than directly releasing individuals with disrupted target alleles.

Aside from the type of genetic control system, ecological context can also have a profound impact on the efficiency of suppression. For example, overcompensation may happen when some larvae die at an early stage and leave resources to other offspring, leading to even more adults in the next generation due to reduced competition [75]. Such resilience varies in different species and regions, influencing several pest control programs [76]. It is therefore important to learn how different strategies respond to varying density dependence.

While a substantial body of previous work has focused on assessing self-limiting genetic control strategies with simulations or mathematical models [16, 68–70, 74, 77–81], most of them modelled ideal systems or had specific ecologies, which creates difficulties when comparing different strategies. In this study, we compared various types of genetic control strategies for population suppression involving repeated releases, focusing mostly on self-limiting strategies and also including some new designs. We also systematically explored the impact of ecological parameters on some of these systems. Finally, we introduce “constant-population genetic load” to evaluate the power of different systems, potentially predicting suppression results and allowing for easy comparisons of different systems and release strategies independent of ecology.

## 2. Methods

### 2.1. Genetic Control Strategies

Our genetic control strategies fall into three classes: modified alleles, gene drives and gene disruptors (Figure 1).

**Figure 1.**
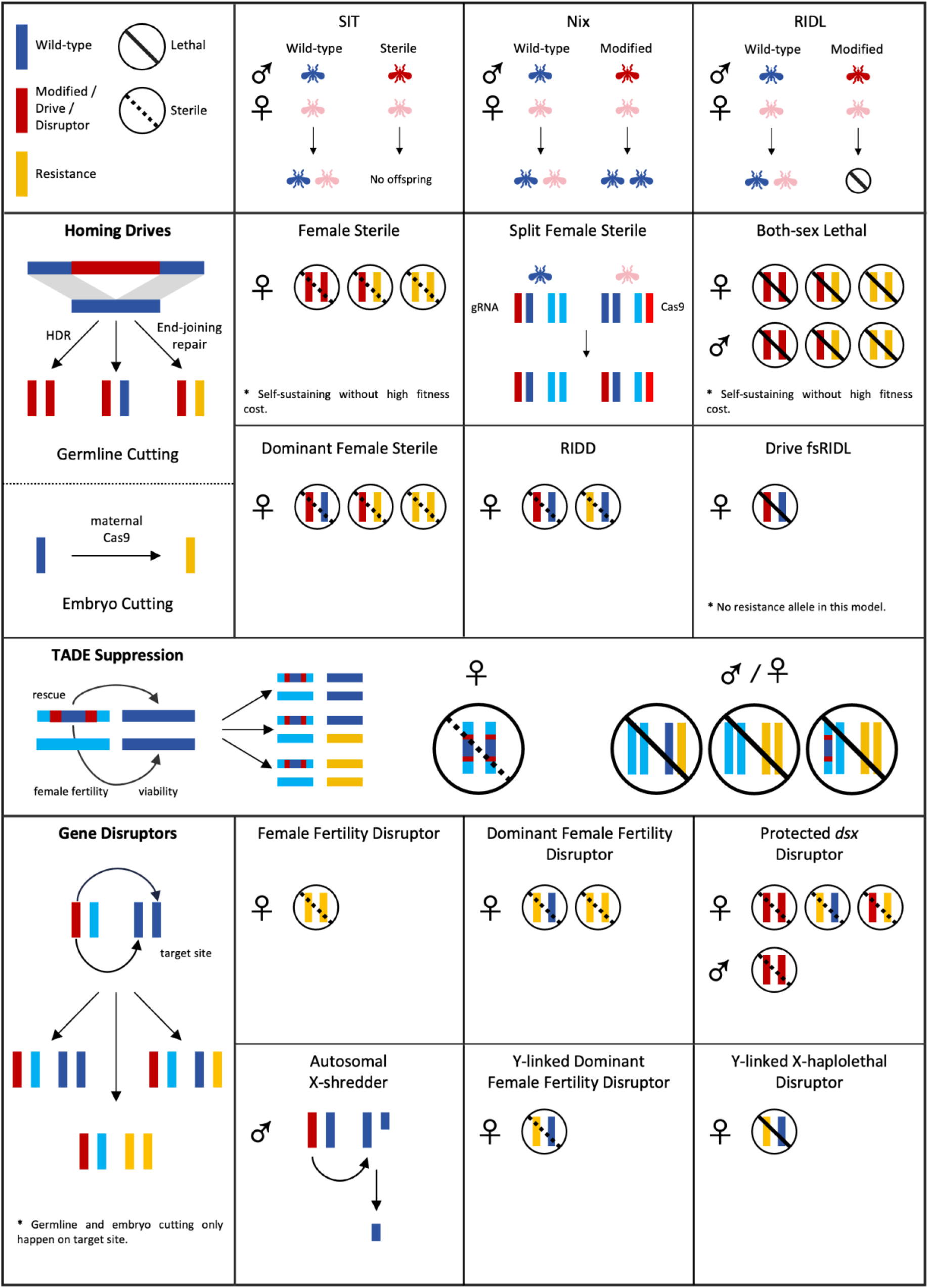
Overview of systems and mechanisms. Each system’s mechanism for population suppression. RIDL can be early or late acting, affecting larval competition. For gene drive heterozygotes, Cas9 causes cleavage at target sites in germline cells, leading to drive conversion or nonfunctional resistance allele formation (only the latter can happen in TADE suppression drive and gene disruptors). Maternal deposition of Cas9 and gRNA may also form resistance alleles in the early embryo. Genetic control systems may result in sterile, non-viable, or sex-biased offspring, depending on their mechanisms.

#### 2.1.1. Modified allele strategies

We assess several types of genetic population suppression strategies. For SIT, sterile males (irradiated or genetic engineered) are released into the population. When females mate with these males, no viable offspring will be generated.

RIDL elements contain a dominant lethal gene that can be suppressed in lab environment (e.g. with food additives such as tetracycline). Male RIDL homozygotes are released into the population, and all of their offspring (or just female offspring for fsRIDL) will die before reaching adulthood but still participate in larval competition. For early RIDL and fsRIDL, offspring are nonviable before the larval stage and don’t contribute to the competition. Therefore, SIT and early RIDL have the same performance in our simulations since released males won’t have any viable offspring in both situations. In real-world conditions, SIT could potentially have worse performance, but this would depend on the type of sterilization and mating system.

Nix is a male-determination factor in *Aedes* mosquitoes, and stable expression of *Nix* will convert females into fertile males [82, 83]. Wild-type males have one copy of *Nix*. Released males have two *Nix* copies on another pair of autosomes, but only half of them have a natural *Nix* (we also considered releases where all males have a natural copy of *Nix*). This line needs to be maintained by using repressible *nix*, similar to RIDL systems. Offspring with at least one copy of *Nix* will become fertile males. Note that *Ae. aegypti* males converted from females through expression of *Nix* alone are flightless, because they lack expression of the *myo-sex* gene responsible for male flight, which is critical for mating in the wild [82]. It may be possible to co-express of *Nix* and *myo-sex* to restore flight in males. However, in *Ae. Albopictus*, *Nix* masculinizes females while keeping their full flight capability [84]. Thus, we model *Nix* males as fully fertile and competitive. The system works to suppress pest populations by biasing the sex ratio.

#### 2.1.2. Gene drive strategies

CRISPR-based drives represent a well-studied and promising class of gene drives. Cas9 and gRNA elements are usually combined with a germline-specific promoter, and ideally, Cas9 only cut the target site in germline cells. However, somatic expression has been a challenge for suppression drives because they usually target a necessary gene related to fertility or viability without rescue. Disruption of these genes in somatic cells leads to fitness costs (30% for drive heterozygotes by default), which can affect females or both sexes, depending on the type of gene drive [85].

For CRISPR homing drives, homology-directed repair after germline cleavage converts the wild type allele into a drive allele in germline cells of drive heterozygotes at a rate equal to the drive conversion efficiency, forming a super-mendelian inheritance pattern. In some cases, end-joining repair takes place and forms resistance alleles, which prevent future drive conversion due to sequence changes. Functional resistance alleles (R1 alleles) will easily outcompete drive alleles but can be avoided by use of multiple gRNAs and conserved target sites [39, 54, 86, 87]. In our simulations, only non-functional resistance alleles (R2 alleles) were included. The total germline cut rate was set to 1 by default, which means all wild type alleles in germline cells of drive heterozygotes are converted into drive or R2 alleles. If the mother was a drive carrier, half of wild type alleles in the embryos would also be converted into R2 alleles due to maternal deposition of Cas9 and gRNA.

Here we modelled mainly three categories of CRISPR drives: 1) self-sustaining homing suppression drives, 2) self-limiting homing suppression drives, and 3) confined toxin-antidote dominant embryo (TADE) suppression drive.

The first self-sustaining drive we considered is the female sterile homing drive. It targets a haplosufficient female fertility gene, so females without any wild-type alleles will be sterile. A well-studied target for such a drive is the *doublesex* (*dsx*) gene, which regulates sex differentiation in insects. Males and females exhibit distinct splicing mechanisms for *dsx* transcripts [88], and targeting the female-specific exon can block functional protein formation and cause recessive female sterility. Another self-sustaining drive is the both-sex lethal drive, which targets a haplosufficient viability gene. Offspring without wild type alleles are non-viable and don’t contribute to larval competition.

Female sterile homing drive can be converted to a self-limiting drive by separating gRNA and Cas9 elements onto different chromosomes. The gRNAs sit in the female fertility gene and causes recessive female sterility, while the Cas9 is at a different genomic location (the drive will functional similarly if these elements are reversed). Drive conversion only occurs when both gRNA and Cas9 are present in the same cell, and this will only increase the frequency of the gRNAs in the female fertility gene. The Cas9, however, is not able to increase its frequency and will be lost when in sterile females, which makes this drive self-limiting. Another example of self-limiting drives is the dominant female sterile homing drive. It also targets a female fertility gene, but also has somatic expression that disrupts wild-type alleles, thus causing dominant female sterility in individuals that are initially drive/wild-type heterozygotes. More recently, RIDD (release of insects carrying a dominant-sterile drive) was developed, in which both drive and resistance alleles cause dominant sterility in females by using multiple gRNAs targeting to the female exon of *dsx*. The last kind of self-limiting drive we explored is drive-fsRIDL. It is similar to fsRIDL, but linked with a CRISPR/Cas9 element, allowing it to spread rapidly while still being self-limiting due to dominant female lethality.

All drives described above are homing drives and rely on homology-directed repair to increase their frequencies. Toxin-antidote dominant embryo (TADE) suppression drive, however, has a distinct mechanism. The drive element sits in a female fertility gene and causes recessive sterility. Meanwhile, the gRNA targets a distant haplolethal gene, which results in dominant lethal disrupted alleles, which can be rescued by a recoded copy within the drive. If an offspring bears more disrupted haplolethal alleles than drive alleles, it becomes non-viable, and females with two drive alleles are sterile. TADE increases its frequency by eliminating wild-type alleles, but requires an introduction ratio above certain threshold if there are any fitness costs or resistance allele formation in the early embryo due to maternal deposition. Therefore, it is confined and will not invade neighbor populations unless migration is above a critical frequency.

#### 2.1.3. Gene disruptor strategies

Gene disruptors are less powerful than gene drive systems because they cannot bias their inheritance, so homozygous release are often required to achieve suppression. These systems are normal alleles with CRISPR/Cas9 or other nucleases that target and disrupt specific genes required for fertility or viability. No rescue is carried by the disruptor. To maintain a stable homozygous line, conditional Cas9 expression is needed. Small molecules, heat shock and optical strategies have been reported to generate conditionally controlled Cas9 cutting [89]. These allow consideration of homozygous releases because homozygotes can be generated prior to any Cas9 cleavage, which would only take place in released organisms.

We started with gene disruptor strategies similar to previously described drives, including both recessive and dominant female fertility disruptors. Another disruptor we considered is the autosomal X-shredder. In this case, any repeated sequence can be targeted even if not within a gene, and the goal for males carrying this shredder to have fewer or no female offspring. The sex ratio of the population is thus biased toward males and facilitates suppression.

Next, we propose a protected *dsx* disruptor, which sits in an early both-sex exon of *dsx* gene and targets its female-specific exon as in the RIDD system above. The disruptor itself is recessive both-sex sterile, and the disrupted target site is dominant female sterile. A similar system “protected dominant negative editor” was also recently proposed with some variations [78].

*Transformer* (*tra*) gene and its paralogs are key switches of sex determination in multiple insects. Knockdown of *tra* causes masculinization and converts XX individuals into intersex individauls or fertile males in *Lucilia cuprina* [90], *Bactrocera dorsalis*[91], *C. capitata* [92] and *Musca domestica* [93]. We modelled a population suppression system targeting *tra*, where individuals with no functional *tra* copy became fertile males.

Finally, we modelled two Y-linked disruptors that only disrupt target genes in the male germline. One is the Y-linked female fertility disruptor, forming recessive female sterile alleles, and the other targets a haplolethal gene on the X chromosome, which leads to dominant lethality in female offspring (males will never receive the dominant lethal allele).

#### 2.1.4. Fitness costs

Fitness costs refer to reduced reproductivity caused by the suppression systems. When targeting essential genes, heterozygotes have a lower fitness (*F* = 0.7 by default) due to somatic expression and disruption of target genes. Fitness costs may apply to males and females or just females, depending on the target gene, leading to less male mating success or fewer offspring per female.

For female fertility drives, female drive heterozygotes have a fitness value equal to *F*. For both-sex viability drive, drive heterozygotes of both sexes are influenced. For TADE suppression drive, if a drive carrier has the same number of drive alleles and resistance alleles, its fitness equals *F*. If it has more drive alleles, it endures a lower fitness of *F^2^*. If it has more functional gene copies, it has a fitness of 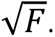. Similar equations are used for fitness costs in gene disruptors. No fitness cost was imposed on SIT, RIDL, fsRIDL and drive fsRIDL to perform a better base line. *Nix*, autosomal X-shredder, *Tra* disruptor and Y-linked disruptors had no fitness cost in our models due to their special mechanisms. All these are likely to have small fitness costs realistically, but in our default parameters, we only model the moderate fitness costs from somatic nuclease expression and other aspects of essential gene targeting with high expression that are particularly difficult to avoid.

In some simulations, an additional fitness cost was imposed on all released males that reduced their mating success. This did not apply to native-born transgenic males.

### 2.2. Panmictic model

We use SLiM software (version 4.2.2) [94] to develop individual-based panmictic models with overlapping generations that approximates relatively short-lived insects that undergo larval competition (See default parameters in Table S1). We start the simulation with a wild-type population with *K* = 100,000 by default. The age structure of this population is 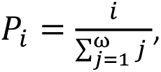 where *i* is the age of individual, and *ω* is the maximum lifespan, which is 6 in our models. The survival rate of adults (age > 0) is age related: 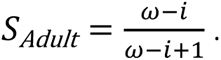

After equilibrating for 10 weeks, we release 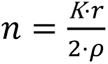 transgenic age 1 males (heterozygotes for drives, homozygotes for modified alleles and disruptors except where noted) into the population each week. Here, *ρ* is the average lifespan. *r* is the release ratio of transgenic males to wild-type males in the population per generation (so fewer males are released each week - the generation time is 2.67 weeks). We also used single releases with *r =* 1 for the self-sustaining female sterile homing drive, *r =* 3 for both-sex lethal homing drive and *r =* 10 for TADE suppression drive. Female fecundity and male mating attractiveness are decided by the fitness value (*p*), which is determined from an individual’s genotype. Every week, each fertile adult female individual randomly selects a male, and will then mate with the male with a success rate equal to the male’s fitness. If this check is failed, the female will select another potential candidate. Each female has a maximum of 10 chances to mate; if none of them succeed or if the selected male is infertile, she will not produce offspring this week. The number of offspring produced is calculated based on the female’s fitness and is drawn from a Poisson distribution with an average of 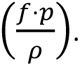 Based on their genotype, nonviable offspring are then removed from the population. All offspring then undergo larval competition. The competition ratio (relative competition compared to the population at normal equilibrium without any transgenic individuals) of larva (*λ*) is determined by the number of viable offspring (*α*): 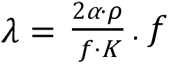 is the average offspring number from a single wild-type female. The actual survival rate of larva is affected by the competition ratio, shape of the density growth curve, and the low-density growth rate ( *β* ). Our models use a linear density growth curve unless otherwise indicated: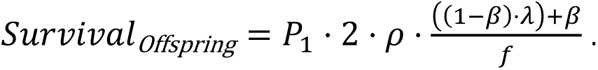

We also explored the influence of a concave curve 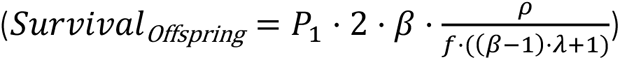 and a convex curve 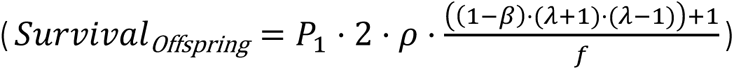 density growth rate. To avoid excessive fluctuation as low-density growth rate increased, we modified the linear and convex curves for some simulations. They were applied only when competition was below the normal level, and if offspring number was boosted above that threshold due to overcompensation, the concave curve would be used to stabilize the population. To perhaps simulate real-world ecological dynamics in many scenarios, a combination of concave and linear curves was used in some simulations. A density character α, ranging from 0 to 1, was employed to control the shape of the mixed curve and to represent the proportion contributed by the linear component (Figure S1).

Numbers of adult fertile females are recorded each week. If there’s no fertile female in the population before week 267 (within 100 generations), the simulation ends in successful population elimination. If suppression fails, the average number of fertile females from the latter half of simulation (week 133-267) is used to show the remaining population size. The minimum release ratio for elimination is determined by binary search until five out of ten simulations achieved elimination, which was considered to be the critical release threshold.

### 2.3. Constant-population genetic load

Genetic load refers to the relative fitness of a whole population compared to a similarly sized wild-type population. Reduced fitness can be caused by mutations or introduction of deleterious genes [95]. For gene drives, it is often defined as the loss of reproductivity of the population. Genetic load is a direct measurement of gene drive power and an accurate predictor of its performance in panmictic models. For self-sustaining gene drives, genetic load for a drive is usually recorded when it reaches its equilibrium frequency. It serves as a good measure of the suppressive power of a drive because it is not affected by species-specific and ecological parameters such as the low-density growth rate.

For systems that involve repeated releases, the dynamics of genetic load is more complex due to increasing relative release ratio as the wild-type population declines. Equilibrium could be reached at a variable population size, which can change the genetic load, even for the same drive and release size, due to ecological factors. We thus sought to report a consistent parameter to allow for better comparisons of repeated release strategies that was based on transgenic system performance and release level. To do this, we propose the concept of “constant-population genetic load”.

If the population size is artificially kept constant during repeated releases, the relative release ratio remains the same, and the relative genotypes will eventually reach an equilibrium. We achieve this by calculating the number of viable offspring based on the drive mechanism, number of fertile females, and proportion of different genotypes. The average offspring number was then adjusted to keep viable juvenile production at the natural level. For most systems (though not late-acting RIDL and several that cause sex-biases), it also means that the native female population is kept around the original level. We then measure the average constant-population genetic load of 50 weeks after the system reaches equilibrium. Equations for each strategy can be found in our models.

### 2.4. Data collection

All simulations were carried out on the High-Performance Computing Platform of the Center of Life Science at Peking University. Data were analyzed and visualized using Python. All models, data, and scripts are available on GitHub (https://github.com/Hanyue22/Assessment-of-Self-limiting-Systems).

## 3. Results

### 3.1 SIT, RIDL, and self-limiting gene drive

To broadly compare different types of population suppression systems with repeated releases, we first evaluated the performance of several self-limiting systems (Figure *2*). SIT eliminated the population when the release ratio was above 4.24, representing the number of transgenic males that are released per generation for each male in the wild-type population at its normal carrying capacity. Note that if SIT releases were insufficient to eliminate the population, the final equilibrium size was boosted above normal carrying due to reduced larval competition and subsequent higher survival of offspring. This was due to our choice of a linear density growth curve and does not occur with a concave Beverton-Holt curve. Density growth curves likely differ between species, but these effects are consistent with previous reports using linear density growth curves [16] and some (though not all) field observations of mosquitoes [76]. Note also that we applied no fitness costs for SIT, which means the transgenic male’s mating success rate was equal to wild type males. This may be consistent with laboratory performance of genetically engineered sterile males, but SIT males generated with methods such as irradiation will likely have a higher fitness cost [96] and requires a larger release size to achieve suppression [97, 98].

**Figure 2.**
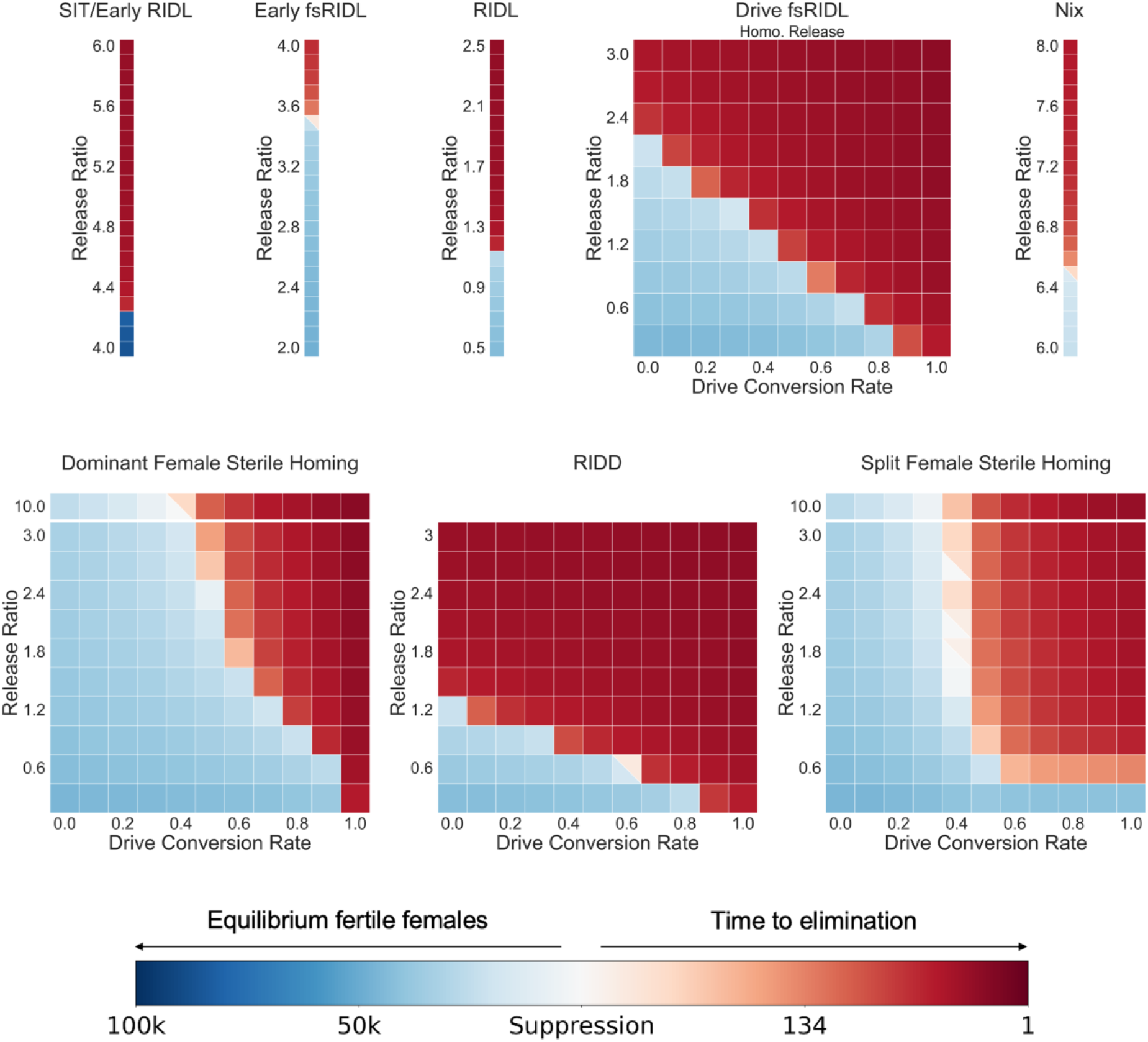
Performance of self-limiting systems. Transgenic males with varying suppression systems were released each week at the per-generation release ratios shown into a wild-type population with a carrying capacity of 100,000. With default performance parameters and varying drive conversion and release ratios, we recorded the time to population elimination or the average number of fertile females after the system reached equilibrium. Squares with two colors indicate that both success and failure happened in simulations. Each point had 10 replicates.

Early acting RIDL is equivalent to SIT, but early fsRIDL is more efficient. Not only are insects easier to rear (the system itself will remove females prior to release), but male offspring, which do not directly contribute to reproduction, will still contribute to larval competition and will still pass on female-lethal alleles to half of their offspring. Thus, the critical release ratio falls from 4.24 to 3.51 (Figure *2*). Late-acting RIDL allows larvae carrying a lethal allele to contribute to competition, enhancing the suppressive effect and substantially reducing the critical release ratio to 1.12. Both-sex RIDL performed better than fsRIDL (with a critical release ratio of 2.17 - see “Drive fsRIDL” in Figure 2 with zero drive conversion) under default settings because wild-type alleles were eliminated from all offspring of released males in RIDL but not fsRIDL. For fsRIDL, released males produce surviving male heterozygotes, which can later mate with females and have half of their daughters survive. In contrast, offspring of RIDL homozygotes all die before reaching adulthood, sacrificing some RIDL alleles in male heterozygotes to more quickly remove wild-type alleles. Thus, the relative release ratio increased. This advantage, however, was not consistent under all ecological parameters (see section 3.6).

The suppression power of fsRIDL was greatly enhanced by combination with homing gene drive (Figure *2*) [77]. Here, we assumed no resistance alleles. The critical release ratio linearly declined as drive conversion rate increased, becoming equivalent to RIDL at 50% drive conversion while retaining the advantage of more efficient rearing. At 100% drive conversion (and no fitness costs), the system remains at the release frequency, so any level of repeated release can eliminate the population. Resistance alleles would decrease drive fsRIDL’s efficiency, but could be largely avoided with an essential target and rescue system in the drive, together with other precautions against functional resistance [77].

*Nix* is a gene responsible for sex determination in several species and has been proposed to be used for genetic control [82–84]. A self-sustaining suppression drive could be developed by linking it to a homing drive, but a self-limiting system could also be developed. By repressing *Nix* expression in a rearing facility, individuals that are homozygous for a *nix* transgene could be generated, half of which would also contain a native copy of *nix*. All progeny of these released males would be males with one or two copies of *nix*. Though this strategy allowed for maximum larval competition, it needed a relatively higher release ratio (6.5) for population elimination (Figure *2*). This is because it does not directly affect female fertility, only converting some to males, which themselves will not have all male offspring if they mate (thus competing with subsequent released males). If combined with genetic or mechanical sex-sorting in a rearing facility, males with three copies of *Nix* could be obtained, reducing the release ratio to 5.47 (Figure S2A). The efficiency of this system could be further improved with additional repressible *Nix* transgene loci, unless expression of even more *Nix* copies caused significant fitness costs.

Dominant female sterile homing drives have been generated when somatic Cas9 expression eliminates wild-type alleles in drive heterozygous females [99]. Released males are drive heterozygous, and performance is heavily dependent on the drive conversion rate. We found that this drive could successfully eliminate the population only when the drive conversion rate was above 0.5, though in this case, the required release ratio could reach much lower than fsRIDL or SIT (Figure *2*). Though it would be difficult to achieve, we also modeled homozygote releases (Figure S2B), which was similar to drive-fsRIDL, but with the addition of nonfunctional resistance allele formation. This significantly increased efficiency when drive conversion was below 0.6 and slightly reduced drive power when the conversion rate exceeded 0.8.

If resistance alleles are dominant sterile along with the drive (the RIDD system [100]), then suppressive power is substantially improved (Figure *2*). This allows male progeny to spread sterility to all daughters even if the daughters don’t receive a drive allele. This efficiency is particularly valuable when the drive itself has a low conversion rate. Unlike normal recessive sterile resistance alleles, which can persist longer and also block drive conversion in males (reducing their benefit), the dominant sterile alleles more heavily contribute to population elimination and are eliminated before blocking drive conversion in males.

Another way to make a self-limiting homing suppression drive is to use a split drive system. With this, we released males that were homozygous for Cas9 and heterozygous for the gRNA allele (the drive) that sat inside and targeted a female fertility gene. Only females carrying both Cas9 and gRNA suffered the default 30% fitness cost from somatic Cas9 expression. Despite being a self-limiting system, it required only half the release size for suppression compared with fsRIDL when drive conversion rate reached 0.5 (Figure *2*), likely because both male and female heterozygotes could continue spreading the drive. Below this threshold, its efficiency declined sharply but still performed better than ideal SIT with a drive conversion rate of 0.4. Time to population elimination shortened as the drive conversion rate rose from 0.6 to 1, but the required release ratio remained unchanged because a certain minimum amount of Cas9 was needed in the population for effective spread of split drive and formation of sufficient numbers of sterile drive homozygotes.

### 3.2 Low efficiency self-sustaining gene drives

We next explored how self-sustaining drive performance could be improved with repeated releases. Though these unconfined drives are powerful, they often will reach a high equilibrium frequency without generating enough suppressive power (gene load, representing the reduction in reproductive capacity of the population) to eliminate the target population. For these systems, the drive conversion rate is the main factor affecting drive efficiency, and female sterile homing drive required a conversion rate above 0.9 to succeed with a single release with our default parameters (Figure *3*A). This could be challenging for many organisms. However, with repeated releases, population elimination could be achieved with only a 40% drive conversion rate. The required release ratio decreased sharply from 40% to 60% drive conversion, reducing the required release sizes to a small level.

**Figure 3.**
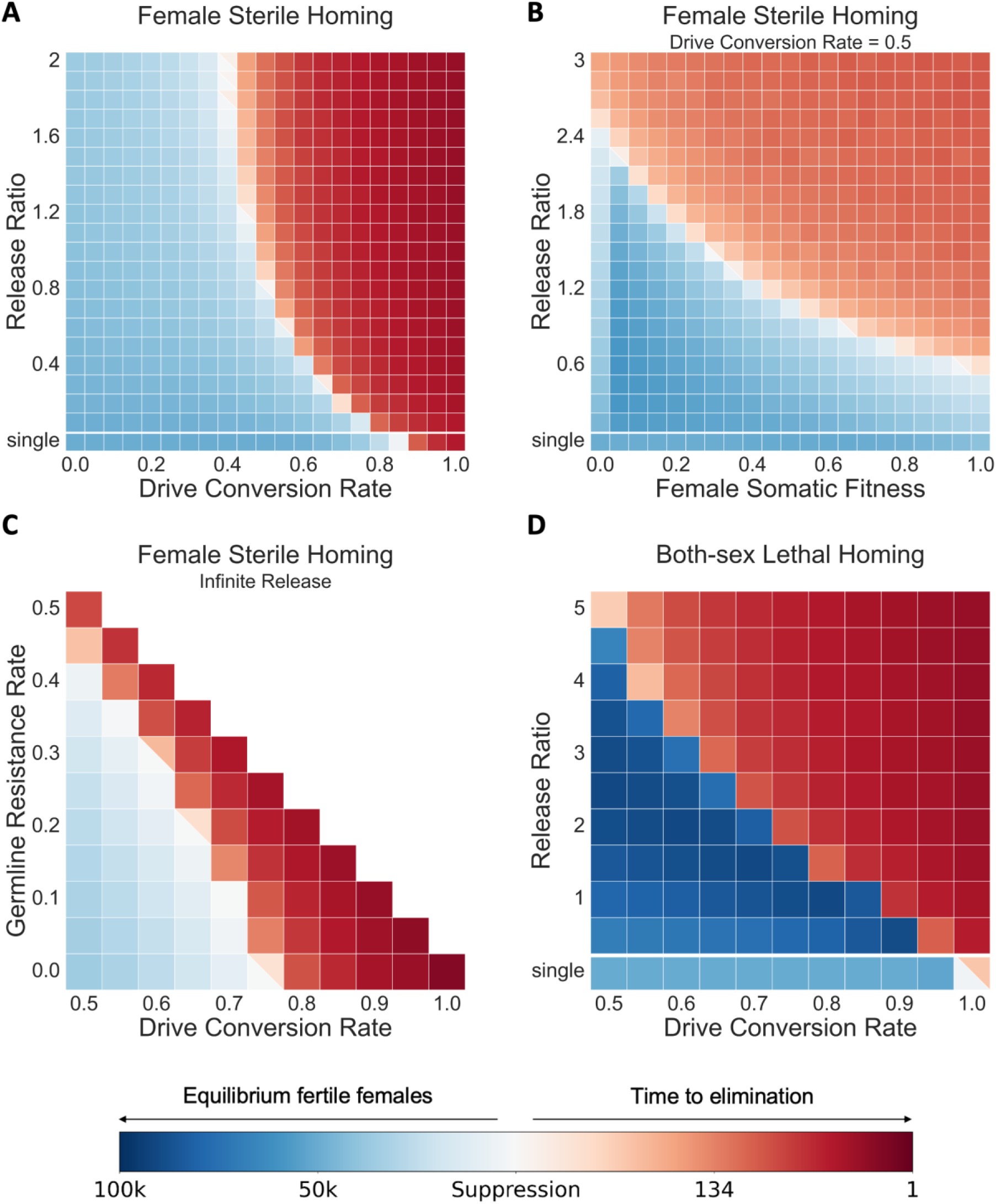
Performance of self-sustaining systems. Transgenic males with varying suppression drives were released each week at the per-generation release ratios shown into a wild-type population with a carrying capacity of 100,000. With default performance except as shown, we recorded the time to population elimination or the average number of fertile females after the system reached equilibrium. Squares with two colors indicate that both success and failure happened in simulations. Each point had 20 replicates for A-B and 10 replicates for C-D.

Another important factor was the drive fitness cost, which is common for this type of drive in heterozygous females if there is any somatic Cas9 expression or requirement for the target gene in the germline. Here we fixed the drive conversion rate at 0.5 and varied the fitness of female drive heterozygotes (Figure *3*B). Low fitness significantly harmed drive efficiency, especially when it was below 0.4, but success was still possible with higher release ratios. With sterile heterozygote females, required release ratio was 4-fold higher compared with a drive causing no fitness cost. Note that when somatic fitness of the drive dropped to 0, female drive heterozygotes were sterile, meaning that there would be fewer fertile females than drives with higher fitness costs. If reduction in fertile females is a primary objective (such as in females in situations where this prevents biting), then dominant drive sterility could be favorable if population elimination is not possible.

We briefly discussed how nonfunctional resistance alleles affected drive efficiency when comparing drive fsRIDL and dominant female sterile homing drive. We also noticed that population elimination with most homing drives (RIDD excepted) becomes impossible with even high release ratios if drive conversion was insufficient. To further investigate these aspects, we conducted an “infinite release” (in which all females were assumed to mate with drive males) while varying both the drive conversion rate and the germline resistance allele formation rate (Figure *3*C). As expected, population elimination could be readily achieved when the drive had high drive conversion, but this was also true with high total germline cut rate (drive conversion and germline resistance allele formation added together), even when drive conversion was moderate. Interestingly, no decrease in drive efficiency caused by resistance allele formation was observed at any drive conversion rate.

Another mechanism for self-sustaining gene drive is to target a haplosufficient but essential gene for both sexes, so individuals with two drive or nonfunctional resistance alleles are nonviable [101, 102]. Though such a drive would have lower genetic load that one targeting female fertility, it could still be a powerful option for suppression with many possible gene targets in different species. We thus explored the performance of a both-sex lethal drive and found that it required a much higher release ratio (Figure *3*D) compared to the female sterile drive. This is partly because male drive heterozygotes would also suffer fitness costs from somatic Cas9 expression (Figure S2C), which we modeled as a reduction in mating competitiveness. Additionally, male homozygotes could not maintain drive alleles, and yet, removal of these males did not reduce population reproductive capacity. This drive also induced overcompensation due to a reduction in viable larva. When we fixed the drive conversion rate at 0.8 and fitness to 0.7, it needed a release ratio of 5 to eliminate the population (Figure S1B). Only very high drive conversion rates allowed for small release sizes.

### 3.3 TADE Suppression drive

TADE suppression drive doesn’t rely on homing to spread, making it slower, but giving the advantage of confinement. When TADE suppression drive has embryo resistance and fitness costs, its introduction threshold increases. Additionally, the main factor influencing its genetic load is the germline cut rate [55], which is easier to improve than drive conversion rate. In our model, TADE suppression drive required at least 70% germline cut rate to succeed (Figure *4*A), either with a single large release above its introduction threshold or with multiple releases. However, larger release sizes did not expand the parameter range for population elimination like in homing suppression drives. However, when the embryo cut rate was high, even a large single release of 10x the population size still resulted in failure (Figure *4*B). In this case, efficiency could be restored using multiple releases. This required larger release sizes but could tolerate somewhat smaller germline cut rates. TADE suppression drive also suffered from overcompensation due to its embryo lethal mechanism, but the effect was much milder because female sterility accounts for most of its suppressive power.

**Figure 4.**
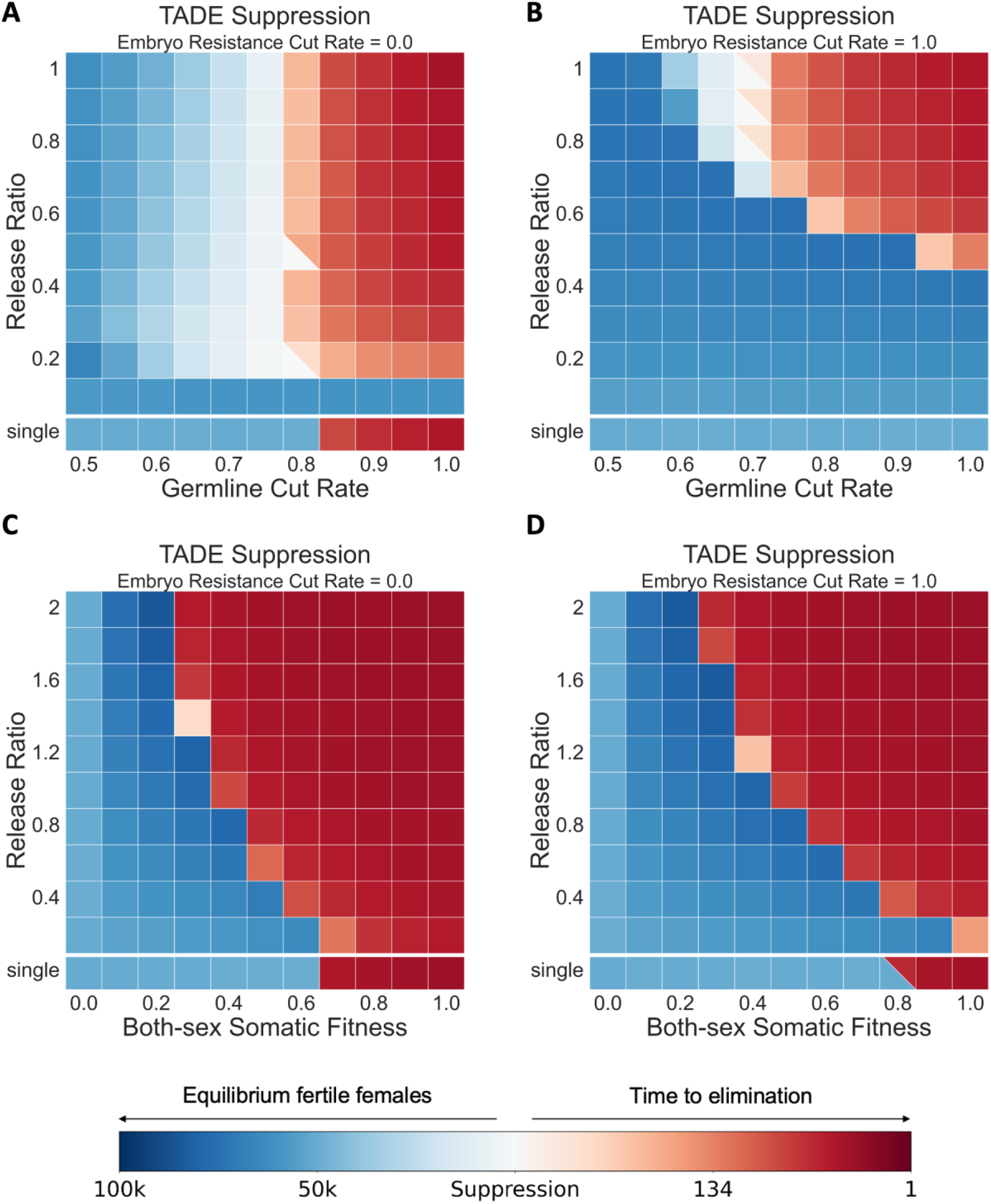
Performance of TADE suppression drive. Transgenic males with TADE suppression drives were released each week at the per-generation release ratios shown into a wild-type population with a carrying capacity of 100,000. With default performance parameters except where indicated, we recorded the time to population elimination or the average number of fertile females after the system reached equilibrium. Squares with two colors indicate that both success and failure happened in simulations. Each point had 10 replicates.

Note that in simulations of single releases for frequency-dependent TADE suppression drive, we set the release ratio as 10. Interestingly, for TADE without embryo cutting, we noticed that a single release performed better than repeated releases, often requiring a smaller total size for more rapid population suppression. At a release ratio of 0.1, success was not even possible because the drive was removed too rapidly from the population. Although the total release amounts were large 0, the introduction ratio in each generation provided a frequency that was far below the drive’s introduction threshold.

We next looked into the effect of fitness cost in TADE suppression drive. Because somatic activity from the drive allele may disrupt the wild-type target gene in somatic cells and couldn’t offer sufficient rescue, both sexes could face high fitness cost. The efficiency of TADE suppression drive was susceptible to this fitness cost (Figure *4*C, Figure *4*D). Even with full germline cut rate, the drive quickly lost its power when somatic fitness dropped below 0.7, or 0.8 when embryo cutting was high. Despite the impact of embryo resistance and fitness costs, TADE suppression drive was relatively powerful compared to SIT and fsRIDL, as long as the germline cut rate was high, a repeated release strategy could cover a deficit in drive fitness, expanding its potential application compared to single release strategies.

### 3.4 Gene disruptors

We define gene disruptors as any construct that targets a certain gene or chromosome without directly providing complete rescue for the disrupted target and without directly increasing their own frequency. We explored the performance of several different self-limiting gene disruptor variants.

First was a simple allele targeting a female fertility gene from a distant site. Such a system, if released in heterozygous form, could not generate sufficient power to eliminate reasonably robust populations. We thus considered homozygous releases, which could be achieved by repressing expression of the construct in the rearing facility. When the germline cutting rate was sufficiently high, the population could be eliminated with a low release ratio of 0.62 (Figure *5*A). Unlike homing drives, where embryo resistance often harmed drive efficiency, cutting in the early embryo reinforced female fertility disruptors. An embryo cut rate of 100% allowed female fertility disruptors to eliminate the population with only 50% germline cutting and an even lower release ratio (Figure *5*B). When we fixed the germline cut rate at 100%, increasing embryo cut rate somewhat decreased the release ratio threshold and duration required for suppression (Figure *5*C). We also modelled severe somatic expression from a female fertility disruptor that led to dominant female sterility and found that it rendered disruptors with a germline cut rate below 0.75 powerful enough to achieve population elimination (Figure S2D), albeit with a higher release size than the version without high somatic activity. In this case, fitness costs from somatic expression reduced population fecundity and favored system efficiency, although it also reduced the persistence of the disruptor.

**Figure 5.**
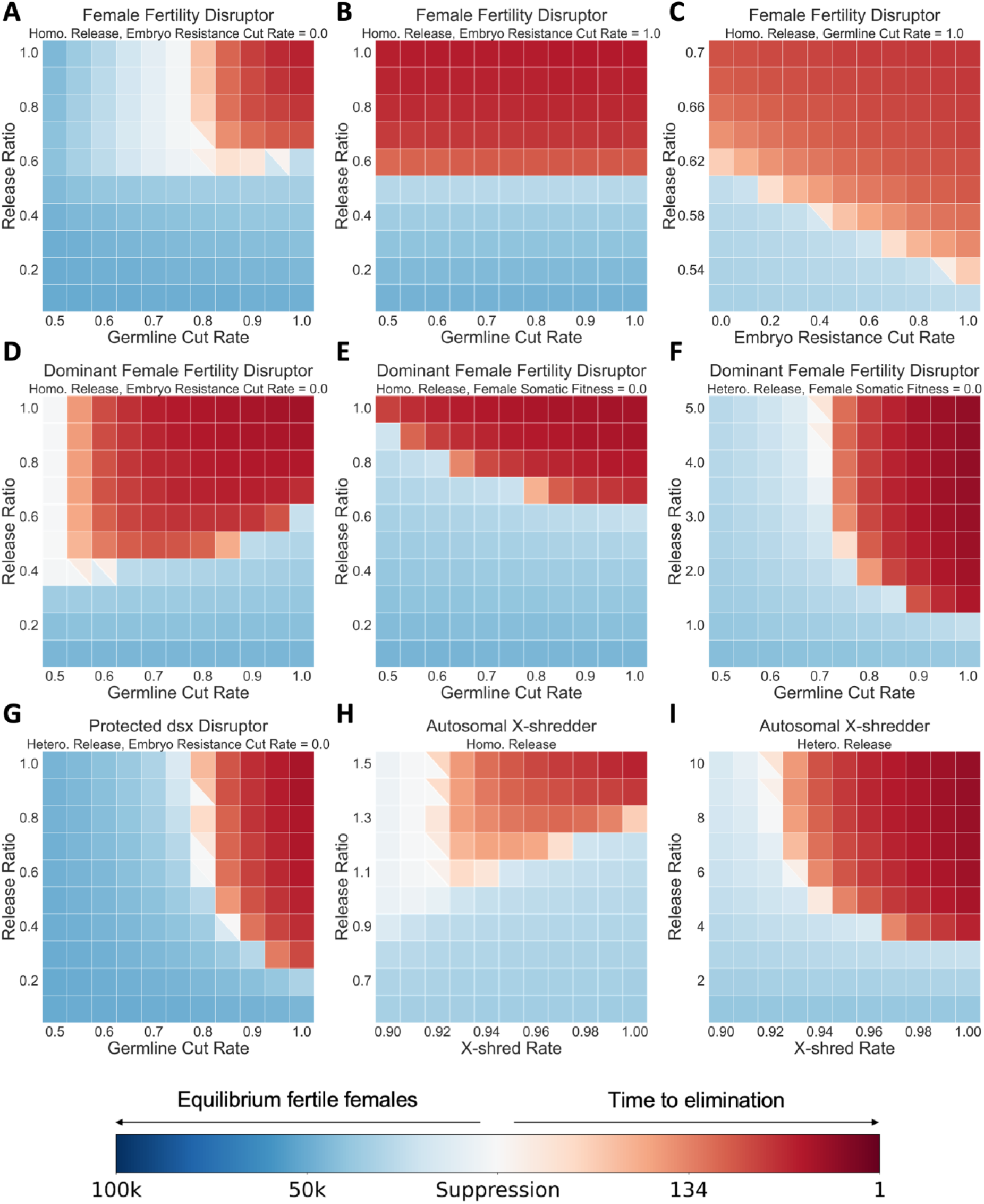
Performance of gene disruptors. Transgenic males with gene disruptors were released each week at the per-generation release ratios shown into a wild-type population with a carrying capacity of 100,000. With default performance parameters except where indicated, we recorded the time to population elimination or the average number of fertile females after the system reached equilibrium. Squares with two colors indicate that both success and failure happened in simulations. Each point had 10 replicates.

Dominant female fertility disruptors form dominant female-sterile alleles in the same way as the RIDD system (Figure *5*D). It was much more powerful than the recessive form, requiring only 55% germline cutting and no embryo cutting to achieve suppression with homozygous releases. It remained effective even if the construct was also dominant female-sterile (Figure *5*E). More efficiency was lost of the release was heterozygous, but it still performed effectively with a high germline cut rate, similar to the RIDD system without any drive conversion (Figure *5*F).

The protected *dsx* disruptor was a distinct system, also based on RIDD in *doublesex*. It targeted a different site on the same chromosome (such as the female exon of *doublesex*), forming dominant female-sterile alleles. However, these alleles would not take effect if on the same chromosome as the disruptor, because the disruptor is placed earlier in the gene, stopping expression and thus preventing dominant sterility. However, drive homozygous females and males would be sterile, lacking any functional copies of *doublesex*. The protected *dsx* disruptor was powerful enough to suppress the population with heterozygous releases (Figure *5*G). When germline cut rate was above 0.8, it allowed population elimination, with a required release ratio of 0.21 when the germline cut rate was 100%. Eliminating fitness cost significantly enhanced its efficiency, lowering the threshold to 0.03 with full germline cut rate (Figure S2E). Unlike most other disruptors, embryo cutting harmed its efficiency, greatly increasing the required release ratio to 0.17 (Figure S2E).

X-shredders have been constructed in *A. gambiae* using both Cas9 and other nucleases [71, 72]. With a repressible system and homozygous release, we found that the system performed fairly well (Figure *5*H). However, heterozygous releases required a much higher release ratio (a release ratio of 3.31 in ideal form) to achieve suppression (Figure *5*I).

Interestingly, for both dominant and recessive female fertility disruptors, as the germline cut rate rose above 0.85, the release ratio threshold for population elimination slightly increased. This unusual pattern was likely caused by resistance/disrupted alleles hindering the spread of disruptor alleles by sterilizing females (and removing disruptor alleles) that would otherwise have been able to further disrupt target alleles in their offspring. For example, in the dominant female fertility disruptor system, when the germline cut rate reached 100%, any female offspring inheriting the disruptor would also inherit a dominant-sterile allele and thus could not pass disruptor or disrupted alleles to the next generation. Compared to a system with only 55% germline cut rate (Figure S3), it generated more resistance alleles at first, but the carrier frequency of the disruptor was eventually limited, reaching a lower equilibrium frequency. A similar pattern emerged in the autosomal X-shredder system with homozygous releases, since few females would be able to inherit the disruptor under high X-shredding rate, and females would tend to enjoy greater reproductive success than males because of the sex bias created by the shredder.

*tra* is a female fertility gene, but in some species, genetic females with *tra* knockout become fertile males. A disruptor targeting *tra* in such a species with an ideal germline cut rate and no embryo resistance could eliminate the population with a homozygous (implying repressibility) release ratio of 1.5 (Figure S2F). Such system might not be applicable to all pest species, but the sex transforming effect allowed resistance alleles to be passed to the next generation, rather than being lost in a female sterile mechanism.

All gene disruptors considered above were located on autosomal chromosomes. Recently, the development of a Y-linked gene editor (YLE) showcases the viability of Y-linked disruptor systems [69]. We explored the performance of a Y-linked dominant female fertility disruptor (Figure S2G), which forms dominant female-sterile alleles at an autosomal site such as *doublesex*. No fitness cost was considered in this model because the disruptor would only be expressed in males. The system was very powerful, with germline cut rates over 0.9 eliminating the population with a release ratio of 0.1. A Y-linked disruptor targeting an X-haplolethal gene was generally as effective (Figure S2H), but took longer because larval competition was reduced (female progeny would be nonviable rather than sterile).

### 3.5 Assessment of suppressive power using constant-population genetic load

For self-sustaining suppression drives, genetic load, representing the reduction in reproductive capacity of the population (suppressive power on a scale from 0 to 1 at the time when the drive reaches equilibrium), is a good measure of long-term system performance, being dependent only on drive characteristics and not ecology, at least in commonly used models. To determine if the drive is powerful enough to eliminate the population, genetic load can be compared to the low-density growth rate. However, it is not possible to easily use genetic load at system equilibrium for assessment suppression strategies involving repeated releases. Performance of genetic control systems with repeated releases are more heavily affected by features of different species and ecological parameters such as the full shape of the density-dependent growth curve. Specifically, genetic load sometimes fails to consistently assess system power due to changes in the relative release ratio. The relative ratio of released males to native-born males will affect the genetic load, but the number of native-born males will change as the population is suppressed, and it will do so in a manner determined by species ecology. Thus, by releasing a constant number of males per generation based on the initial male number (as in our simulations) the genetic load will change as the population size is changed, even if genotype frequencies quickly reach equilibrium. To provide an unchanging genetic load measurement for repeated release systems, we introduce the concept of “constant-population genetic load” to quantify system power. We measure this by artificially adjusting reproduction for all fertile individuals by a particular constant, calculated to maintain the native-born population size. We then allow the system to reach equilibrium during continuous releases and measure the genetic load at this point. Here, we assess the accuracy and consistency of this measurement across multiple suppression strategies.

Constant-population genetic load correlated closely with the performance of different systems (Figure A, Figure S4). Specifically, higher values tended to result in more successful population elimination (compare to other figures) and good correlation with system performance. For example, as the release ratio went up, overcompensation first led to higher average population sizes for both-sex viability homing suppression drive. However, constant-population genetic load results suggested that system power was stably increasing. Constant-population genetic load results for the dominant female sterile disruptor had similar pattern with its actual performance. It suggested that the relatively inferior outcomes when the germline cut rate rose above 0.9 were due to system deficiency rather than environmental impacts.

**Figure 6.**
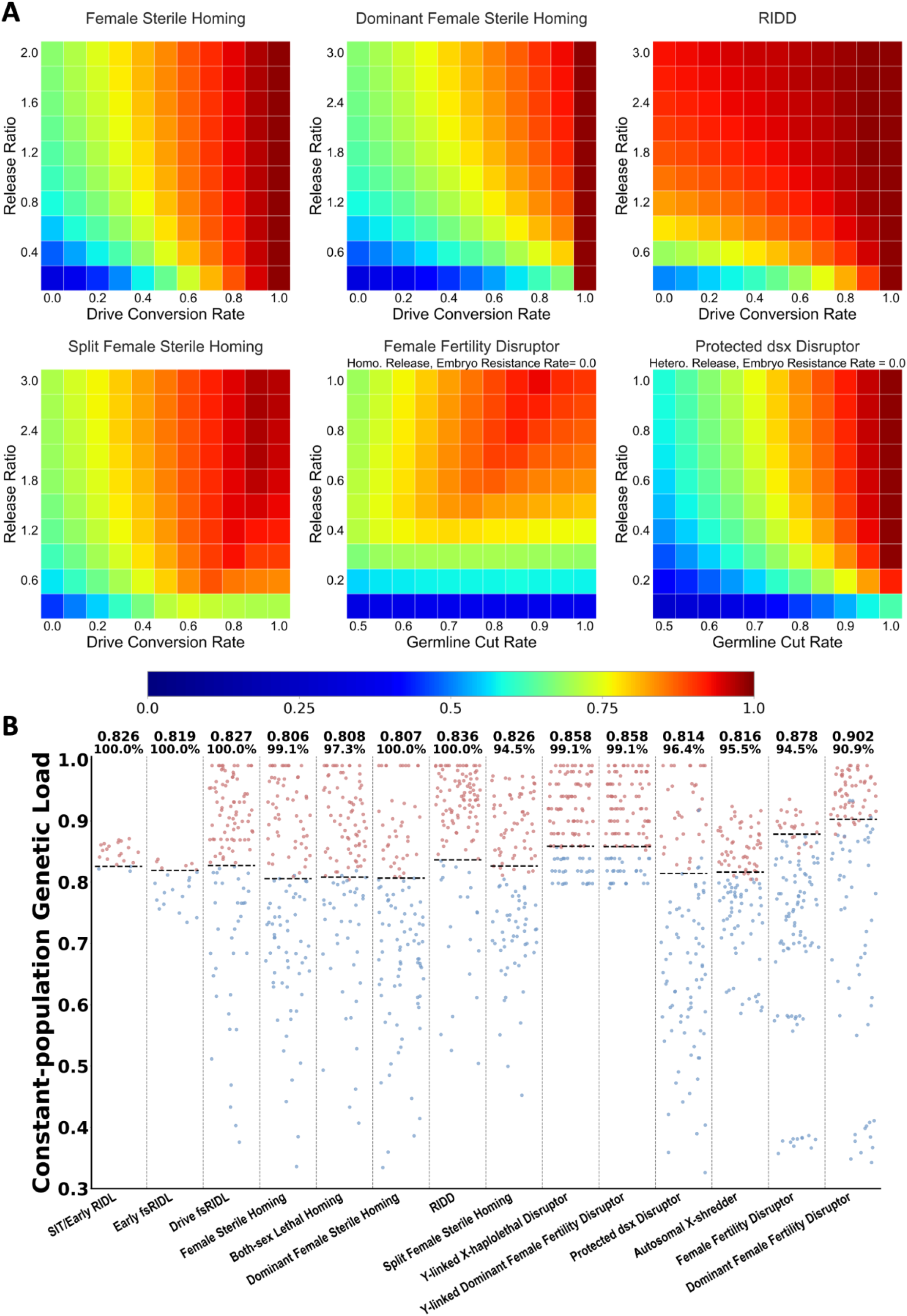
Constant-population genetic load. A. Transgenic males were released each week at the per-generation release ratios shown into a wild-type population, with the population being kept around its carrying capacity of 100,000. With default performance parameters except where indicated, we recorded the average constant-population genetic load for 50 weeks after the system reached equilibrium. Each point had 5 replicates. B. Red and blue dots indicate successful and unsuccessful elimination, respectively, using data obtained from constant-population genetic load simulations and normal simulations. Each dot represents a certain parameter set from our earlier data. Black dashed lines represent the constant-population genetic load at the critical release ratio needed for population elimination. This critical release threshold and the accuracy of using this to predict results in the other points are shown above the graph.

We next explored the relationship between constant-population genetic load and successful elimination. For regular genetic load in self-sustaining systems, the relationship between genetic load and deterministic population elimination is straightforward. The genetic load must be equal or greater than [1 - 1/(low-density growth rate)]. Constant-population genetic load does not provide as simple a relationship because the full shape of the density growth curve (not just the low-density growth rate) and other ecological factors relating to competition will all influence the result. Nevertheless, we still uncovered a strong correlation between elimination outcomes and the constant-population genetic load for many different systems (Figure B). We determined the minimum release ratio for population elimination in different systems with drive conversion rate or germline cut rate of 0.8 (except for the 2 Y-linked disruptors which required a germline cut rate above 0.86 to eliminate the population), and we measured the constant-population genetic load at this critical release ratio. We then compared this to our previous simulations and their associated values for constant-population genetic load. This served as an accurate predictor for population elimination outcomes for SIT, early fsRIDL and most homing drive systems, all at a consistent level when holding ecological parameters constant. However, some systems with more exotic mechanisms (Y-linked disruptors, female fertility disruptors) had their critical constant-population genetic load threshold at a different level, showing that while internally mostly consistent for different drives, this value has less utility comparing certain specific system types than genetic load for self-sustaining gene drives.

We further examined systems where the threshold prediction had a lower accuracy when comparing to our previous simulations (Figure S5). The split female sterile homing drive and autosomal X-shredder were influenced mostly by parameter spaces with binary outcomes (elimination in some simulations and population persistence in others with the same simulation parameters). For the female fertility disruptors (recessive and dominant), accuracy fell near the borders of population elimination. The threshold measured at different performance parameters varied, probably due to the inhibiting effect of high germline cut rate. For RIDL and TADE suppression, the thresholds were difficult to measure in the first place. The late-acting, both-sex lethal mechanism of RIDL hindered the equilibrium state near elimination. For TADE, its frequency-dependent nature and unique mechanism including both-sex lethal and female sterile effects led to rapid population elimination in the constant-population simulations, preventing accurate measurements (this system is usually self-sustaining and best assessed with conventional genetic load).

### 3.6 Impact of density growth curve on population suppression

Ecological factors can greatly affect suppression systems involving repeated releases. In addition to the low-density growth rate, the shape of the density growth curve can play a more important role than for self-sustaining gene drives. We thus explored the performance of different systems under various ecological conditions. The shape of the density growth curve affected population stability and resilience. A “concave”-shaped Beverton-Holt curve resulted in stable population level but low resilience in response to a genetic load. A “convex” curve allowed population size to remain high when facing moderate suppressive power. Our default setting in simulations was a “linear” curve, with intermediate effects. Overcompensation can occur with any curve that is more robust than our concave curve.

We first investigated the impact of low-density growth rate on the four classic suppression systems: RIDL, fsRIDL, early-acting fsRIDL, and SIT (equivalent to early-acting RIDL). We determined the release ratio threshold needed for population elimination with the linear density-dependent growth curve (Figure A). With a low-density growth rate of 2, fsRIDL required the lowest release ratio for population elimination (0.18), outcompeting RIDL since males could still spread the female-specific lethal alleles. When low-density growth rate rose above 4, however, RIDL became more powerful. The two early-acting systems were less efficient throughout entire range. SIT required a higher release ratio when low-density growth rate was below 8 but outperformed early fsRIDL when it was over 10. Overall, the both-sex systems were linearly impacted by low-density growth rate, while female-specific systems were more substantially affected by higher low-density growth rate due to their reduced ability to eliminate wild-type alleles.

**Figure 7.**
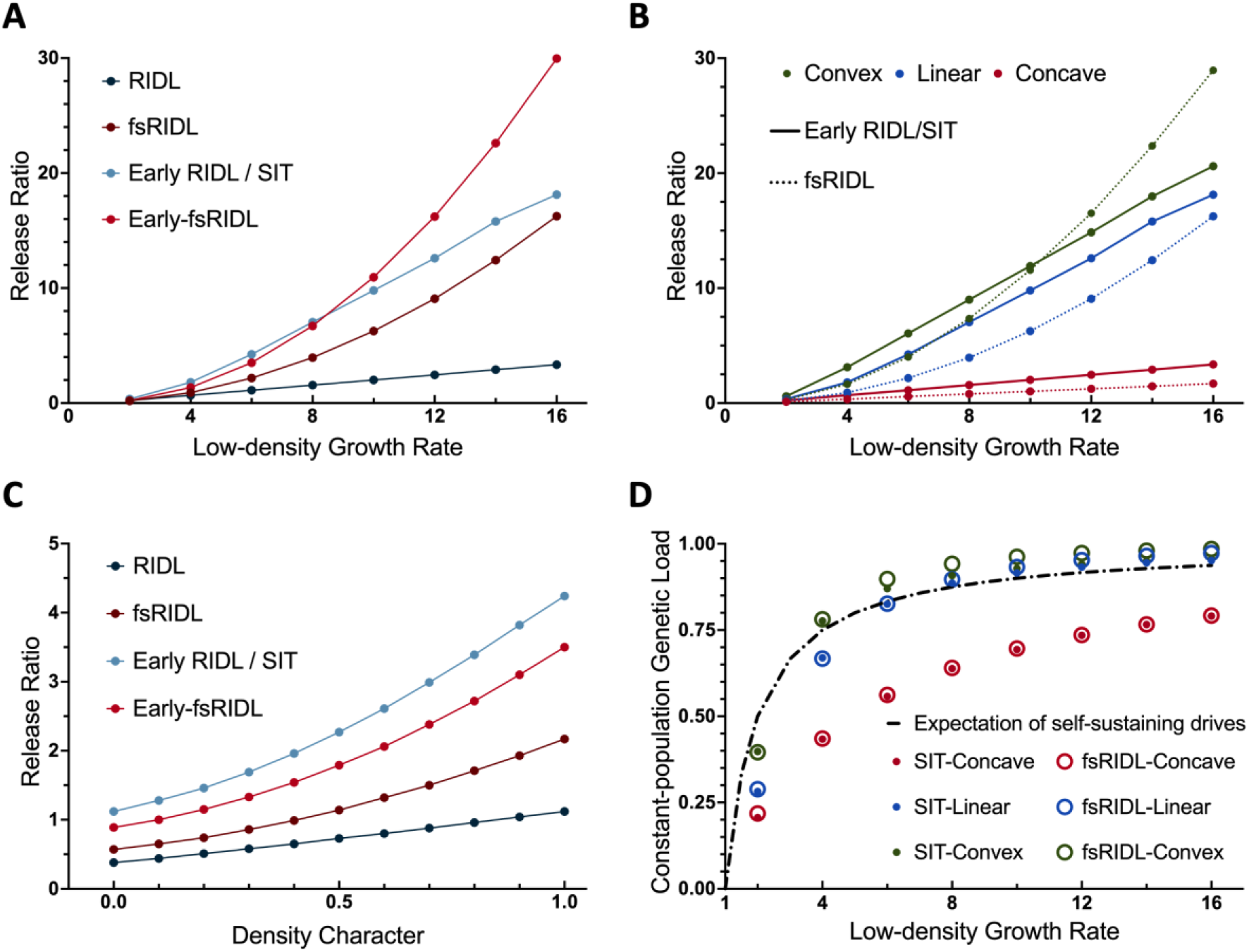
Impact of ecological parameters. Critical release ratio threshold needed for population elimination using several systems are shown for (A) linear density growth curve and varying low-density growth rate, (B) different shaped density growth curves and varying low-density growth rate, and (C) varying density character showing a transition from concave (0) to linear (1) with a low-density growth rate of 6. (D) Constant-population genetic load threshold needed for population elimination with differently shaped of density growth curves with varying low-density growth rates. The black dashed curve represents the genetic load threshold for elimination after a single release of a self-sustaining suppression drive, which is equal to 1-1/β. Each spot had 5 replicates.

We next tested the release ratio threshold of fsRIDL and SIT/early RIDL under different low-density growth rates with all three kinds of curves (Figure B). As expected, higher low-density growth rate increased release sizes. The convex curve required the highest release ratio to achieve elimination, and the concave curve needed the lowest. The fsRIDL system usually performed better, but it was more strongly affected by the low-density growth rate in linear and convex curves. It was only inferior to SIT for the convex curve when low-density growth rate was over 10. This is because the main advantage of fsRIDL over SIT comes from higher larval competition from offspring of released males. If this is less important because population size will remain high until near the point where population elimination is suddenly achieved, then fsRIDL is left with the disadvantage of surviving male progeny of released males carrying wild-type alleles and competing with released males. Because RIDL allows for full competition while removing wild-type alleles rapidly, its critical release ratio increases only linearly as the density curve transitions from concave to linear, while all other systems are more greatly affected (Figure C).

We then measured the corresponding constant-population genetic load at the critical release ratio needed for population elimination for SIT and fsRIDL, comparing them to the theoretical necessary ratio based only on the low-density growth rate (Figure D). Because the predictive threshold of constant-population genetic load was consistent across multiple systems, this result indicated the approximate required intrinsic power for a genetic control system to eliminate the population under different low density growth rate. Consistent with the release ratio result, it was the easiest to suppress the population with a concave curve, and the convex-mix curve required the most suppressive power. We also observed that when low-density growth rate rose above 6, the threshold for linear and convex curves surpassed expectation probably due to overcompensation. However, as the low-density growth rate further increased, thresholds under all circumstances tended to converge to the theoretical expectation.

## 4. Discussion

In this study, we modelled various genetic control systems (Table 1), focusing on self-limiting strategies involving repeated releases of transgenic males. Results suggested that several types of gene drive and gene disruptors could eliminate pest populations substantially more effectively than SIT and RIDL, offering a feasible alternative for population suppression with potentially lower resource investment. To facilitate comparison between the large number of potentially effective systems utilizing repeated releases, we proposed constant-population genetic load to evaluate system power, showing that it is often a useful way to assess these systems irrespective of ecological factors.

**Table 1.** Properties of suppression systems.

|  | Strategy | Ideal Release Ratio | Default Release Ratio | Requires repressibility | Resources used for females | Requires sorting | Requires outcrosses | Sensitive to overcompensation |
| --- | --- | --- | --- | --- | --- | --- | --- | --- |
| Modified Alleles | Genetic SIT/Early-RIDL | 4.24 | 4.24 | No | Yes | Yes | Yes/No | Yes |
|  | RIDL [17] | 1.12 | 1.12 | Yes | Yes | Yes | No | No |
|  | Early-fsRIDL [22] | 3.51 | 3.51 | Yes | No | No | No | No |
|  | fsRIDL [22] | 2.17 | 2.17 | Yes | Yes | No | No | No |
|  | *Nix* [82, 84] * | 5.47/6.5 | 5.47/6.5 | Yes | No | No | No | No |
| Gene Drives | **Self-sustaining drives** |  |  |  |  |  |  |  |
|  | Female sterile homing [39, 72, 103] | <0.01 | 0.05 | No | Yes | Yes | Yes | No |
|  | Both-sex lethal homing [103] | <0.01 | 1.37 | No | Yes | Yes | Yes | Yes |
|  | **Self-limiting drives** |  |  |  |  |  |  |  |
|  | Split female sterile homing [33] | 0.54 | 0.56 | No | Yes | Yes | Yes | No |
|  | Dominant female sterile homing [33] | 0.24 | 0.98 | No | Yes | Yes | Yes | No |
|  | RIDD [100] | 0.03 | 0.35 | No | Yes | No | Yes | No |
|  | Drive-fsRIDL [77] | 0.03 | 0.44 | Yes | No | No | No | No |
|  | **TADE drive** |  |  |  |  |  |  |  |
|  | TADE suppression [55] | <0.01 | 0.17 | No | Yes | Yes | Yes | Partially |
| Gene Disruptors | **Autosomal-linked disruptors** |  |  |  |  |  |  |  |
|  | Female fertility disruptor | 0.45 | 0.62 | Yes | Yes | Yes | No | No |
|  | Dominant female fertility disruptor ** | 0.62/1.07 | 0.62/1.07 | Yes/No | Yes | No | Yes | No |
|  | Protected *dsx* Disruptor [78] | 0.03 | 0.21 | No | Yes | Yes | Yes | No |
|  | X-shredder [71-73, 104] ** | 1.3/3.31 | 1.3/3.31 | Yes/No | No | No/Yes | No/Yes | No |
|  | *Tra* disruptor | 1.46 | 1.49 | Yes | No | No | Yes | No |
|  | **Y-linked disruptors** |  |  |  |  |  |  |  |
|  | Dominant Female fertility disruptor | 0.03 | 0.03 | No | Yes | No | Yes | No |
|  | X-haplolethal disruptor | 0.04 | 0.04 | No | No | No | Yes | No |

Release ratio refers to minimum drop size in each generation for population elimination. 'Ideal' indicates ideal system performance parameters, and 'Default' indicates default parameters. \* Released males with 3/2.5 *Nix* copies. \*\* Homozygous/heterozygous release.

Numerous genetic control strategies have been proposed and verified in many species over the past decades. As researchers start to contemplate field trials for more powerful strategies [51, 79], a central question emerges: which system should we opt for? This can be further deconstructed into four aspects: 1) what is our desired outcome, 2) where do we aim to control the pest, 3) what systems can be built in the target species, and 4) what is the effectiveness of the viable systems? The first question tends to be straightforward, dividing scenarios into population modification or suppression, though the level of desired suppression could vary in some cases. The second question is very important, but also potentially complex. Self-sustaining gene drives and even confined gene drives could potentially be less desired due to a variety of factors.

Once an objective is set, possible strategies may still be limited because not every system can be successfully implemented in a target species due to a lack of genetic knowledge and the absence of universal transgenic technologies applicable to non-model organisms. Even when successful, efficiency can vary. For instance, gene drives with the capability to eliminate population by single release often necessitate a well-characterized target site, reliable gene-editing tools, and promoters with high germline specificity and minimal maternal deposition— all of which may be difficult to meet in some species. According to our modelling results, gene disruptors are usually less powerful than even self-limiting gene drives, relying on repeated releases of homozygous transgenic males in some cases. However, they still outcompete SIT and fsRIDL with high germline cut rate, which is easier to achieve than high drive conversion rate. The self-limiting nature of gene disruptors can mitigate safety concerns, and easy access to conditional/repressible systems (Table 1 “Requires repressibility”) or effective insertion sites on the Y chromosome would make more of them potentially feasible solutions for pest control. With limited options, it is crucial to select a system with relatively better efficacy and feasibility based on simulation outcomes.

Genetic pest control strategies often have eco-evolutionary feedback, which can greatly influence their outcomes, and such impacts are closely correlated with features of targeted species as well as the climate and terrain of targeted areas [105, 106]. The density growth curve, including the low-density growth rate, is a general description of population response to suppressive power. We showed that the shape of density growth curve and low-density growth rate significantly changed the performance of different strategies, suggesting that specific control methods should be applied according to species and environmental conditions. A system outcompeting another in one situation may be a lot weaker under alternative conditions. Therefore, performance of systems and suppressive power required for population elimination in specific species and environments should be fully considered when determining the optimal system if at all possible. This will often not be the case, though due to a lack of ecological data even in many relatively well-studied species. We proposed constant-population genetic load as a measurement of intrinsic system power in repeated releases and explored the suppressive threshold for different systems and conditions. Despite a few exceptions, constant-population genetic load shows good consistency and reliable predictive power for multiple systems, making it a potentially valuable alternative method to consider different systems independently of the specific scenarios that may lack available data.

Economic and labor resources are additional considerations for efficiency that we did not consider here because they would usually vary based on location, pest species, and potentially even specific systems. Especially important for these is the complexity of rearing facilities. For many drives, resources in the final release rearing batches are spent on females, reducing efficiency at this step (Table 1 “Resources used for females”). Mechanical sorting, either by sex or with fluorescence, can also add time, cost, and complexity to rearing (Table 1 “Requires sorting” - note that the tables assumed that intersex females may be released, but not sterile females, but this may sometimes be acceptable). For some systems, it is necessary to rear separate batches of insects and then cross them together (usually requiring sex-sorting) to generate the batch required for deployment, adding an additional consideration (Table 1 “Requires outcrosses”). Some of these can be mitigated in many of these systems by adding genetic complexity to improve efficiency, such as male-killing that takes effect to release batches. All these considerations interact with the costs of deployment per insect and setup/facility maintenance costs to provide a full assessment of resources required for a release program. For example, in our simulations, the release ratio required for RIDL is lower than that for fsRIDL under many conditions. However, fsRIDL can automatically filter out females, thereby reducing costs significantly. Early-acting fsRIDL is less effective than late-acting fsRIDL per released male, but up to twice as many males could be generated for the same amount of food because females will be nonviable before they consume a significant amount of food. Another consideration is when a system has a slightly higher release ratio threshold, but a shorter duration for population elimination, resulting in a smaller total release size. The comprehensive cost of each strategy, including research and development, material preparation, release implementation, as well as long-term management and monitoring, is an important factor for its application prospects.

Achieving a balance between efficiency and controllability has been a substantial challenge in the field of genetic pest control. There are instances where the long-term persistence of genetically modified elements within a population is undesirable, or where we may need the ability to halt the genetic intervention and reverse its impact. Previously proposed ideas including elements that disrupt the drive by releasing another gene drive [107], or simply releasing wild-type individuals to dilute and eliminate drive alleles [108]. Such solutions are costly, and the release of a second kind of transgenic individual may cause further safety concerns, particularly if an initial release was associated with an unexpected problem. A repeated release strategy using a self-limiting system provides an alternative. Compared with SIT and fsRIDL, repeated releases of weak gene drives and disruptors require a much lower release ratio, thereby conserving resources. Meanwhile, in contrast to self-sustaining homing drive that spread from a single introduction, ceasing repeated releases will result in the allele frequency gradually diminishing for self-limiting systems (Figure S6A). This strategy also renders low-cost genetic control feasible in some species where it is difficult to develop a homing system with high drive conversion rate and low fitness costs. Moreover, by selecting appropriate size and duration of repeated releases, this flexible strategy allows complete eradication of target population, maintaining it at low level, or full recovery after a short-term period of suppression (Figure S6B).

Our panmictic models in this study are generalized and may be widely applicable to many different systems where offspring compete. However, other systems are possible. Previous research on spatial models indicates that chasing and long-term persistence of wild-type alleles could happen for self-sustaining suppression drive and hinder the elimination of population [101, 109]. This would be less likely for self-limiting systems released in a limited area, but the nature of migration from outside the area could still drastically change outcomes and increase required release sizes. Progress has also been made in building gene drive models considering interspecies interaction [110], disease transmission [111], and temporal dynamics [112]. Investigating how repeated release strategies and self-limiting systems perform in such scenarios may yield important insights. Constant-population genetic load fills a gap in evaluating the power of repeated release strategies but does not perfectly predict suppression results in some systems. A better understanding of these could result in improved predictions.

It is well understood that released males that were reared in a laboratory environment have reduced mating competitiveness due to a variety of factors. These can include but are not limited to incomplete nutrition, lack of environmental cues, and effects arising from the wait between adult maturation and actual release. Release of eggs instead of adults is possible for many systems, but it remains unclear if this provides greater overall efficiency due to greater egg mortality, which could substantially exceed mortality of naturally laid eggs. We did not model such considerations, which would theoretically increase critical release sizes by a similar factor for all systems considered. However, some systems may have a greater advantage for reducing reared male fitness costs in real-world conditions, potentially giving them an additional advantage.

In summary, we performed a detailed comparison of genetic control strategies for population suppression, including SIT, RIDL, several homing drive variants, TADE suppression drive, and several classes of gene disruptors. Many self-limiting strategies are highly feasible and represent large improvements over current state-of-the art systems, which may be particularly desirable if even more cost-effective suppression gene drives cannot be developed due to technical challenges or cannot be deployed due to ecological, social, or other considerations. Moreover, constant-population genetic load effectively evaluated the power of different repeated release systems, allowing it to serve as a promising predictor of population elimination when comparing strategies. However, ecological factors and rearing economics should also be considered when determining the most promising approach in specific scenarios.

## Acknowledgements

Thanks to the High-Performance Computing Platform of the Center for Life Science at Peking University for assistance with cluster-based data collection. This study was supported by the Center for Life Sciences, as well as the grants from the National Science Foundation of China (32270672) and the Beijing National Science Foundation (Q2023041).

## Supplemental Information

**Table S1.**
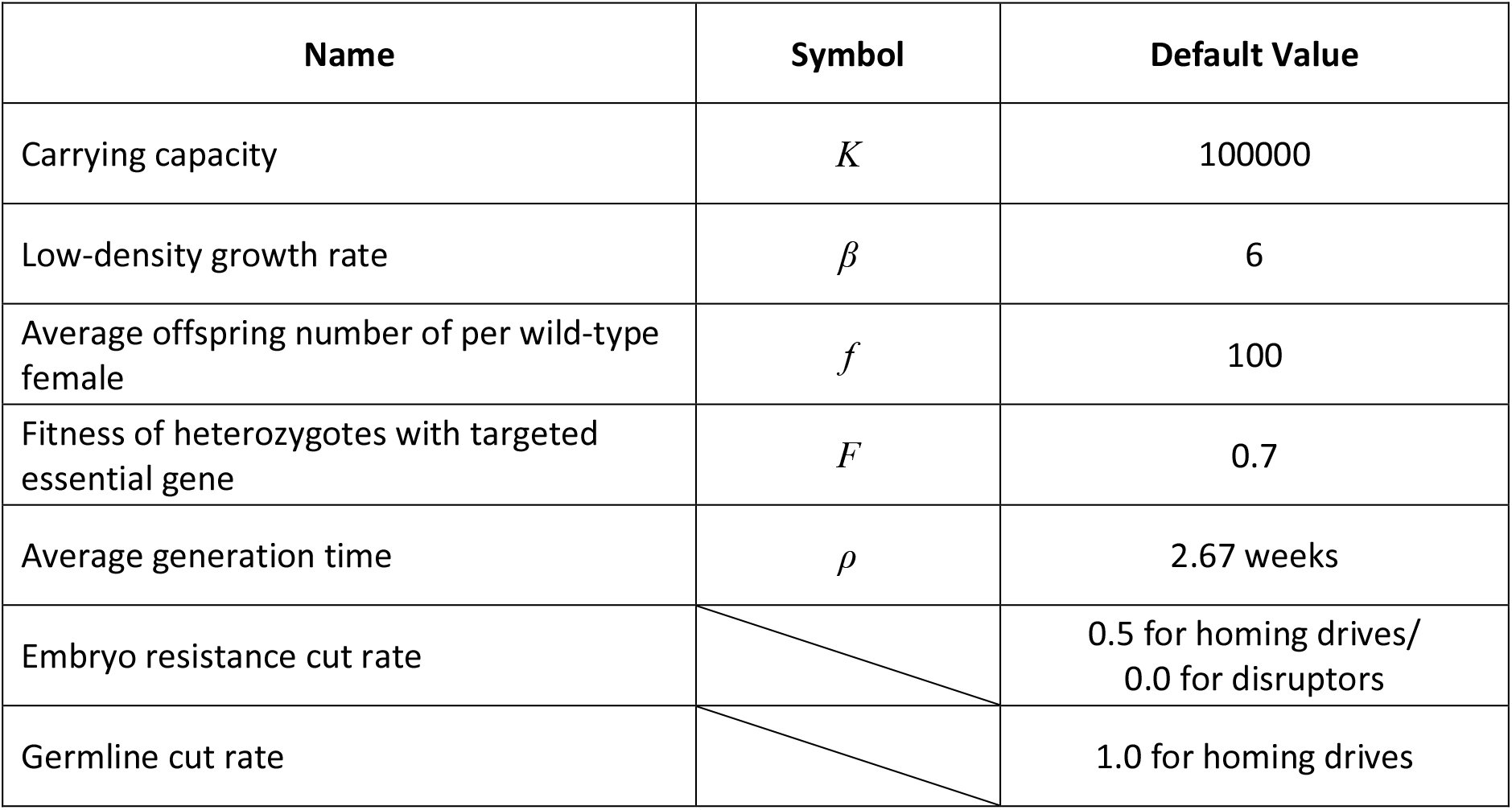
Default parameters for SLiM models.

**Figure S1.**
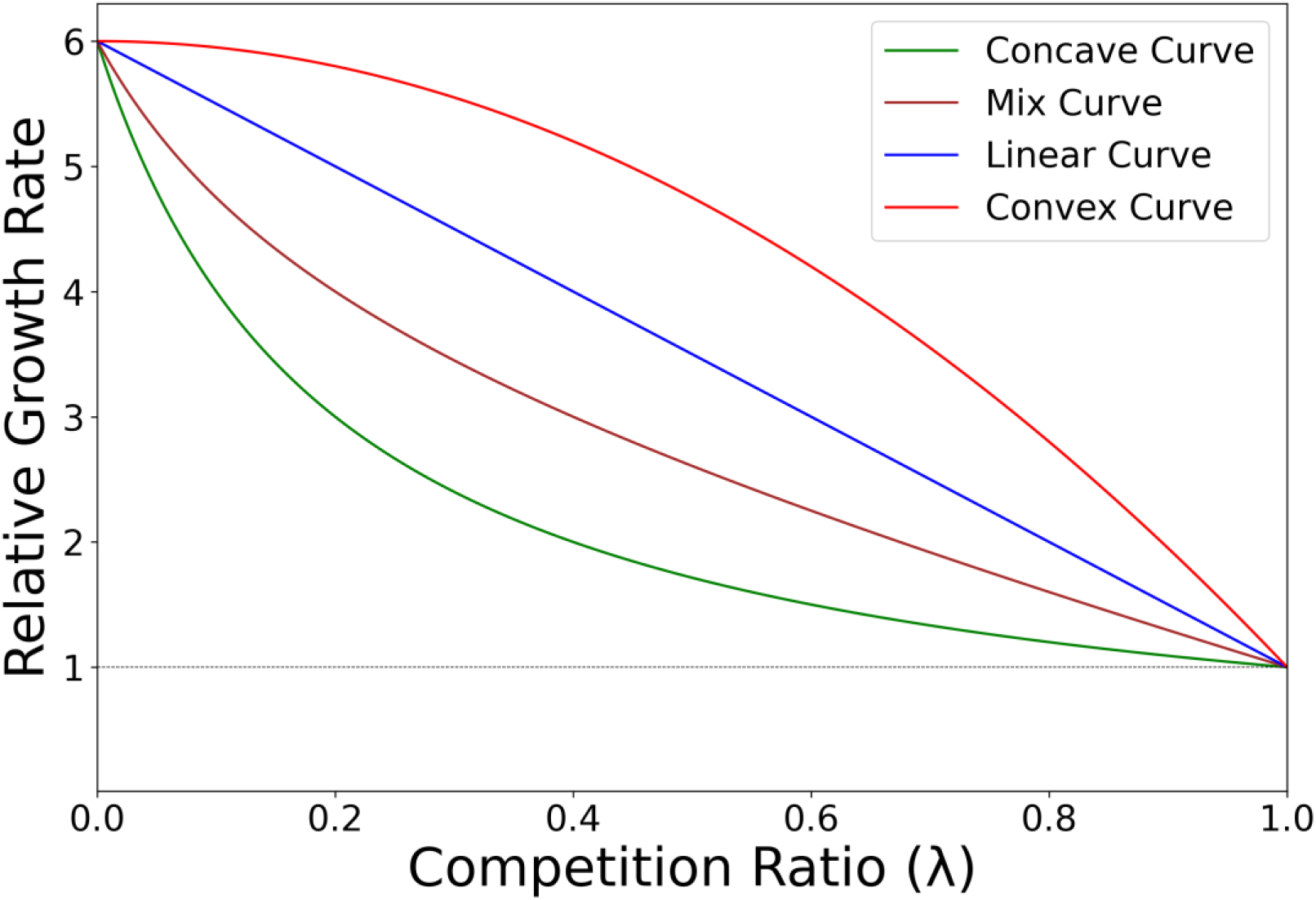
Shapes of density-dependent growth curves. The figure shows the shapes of the concave, linear, convex, and mix curve when the density character is 0.5. The figure was plotted with a default low-density growth rate of 6.

**Figure S2.**
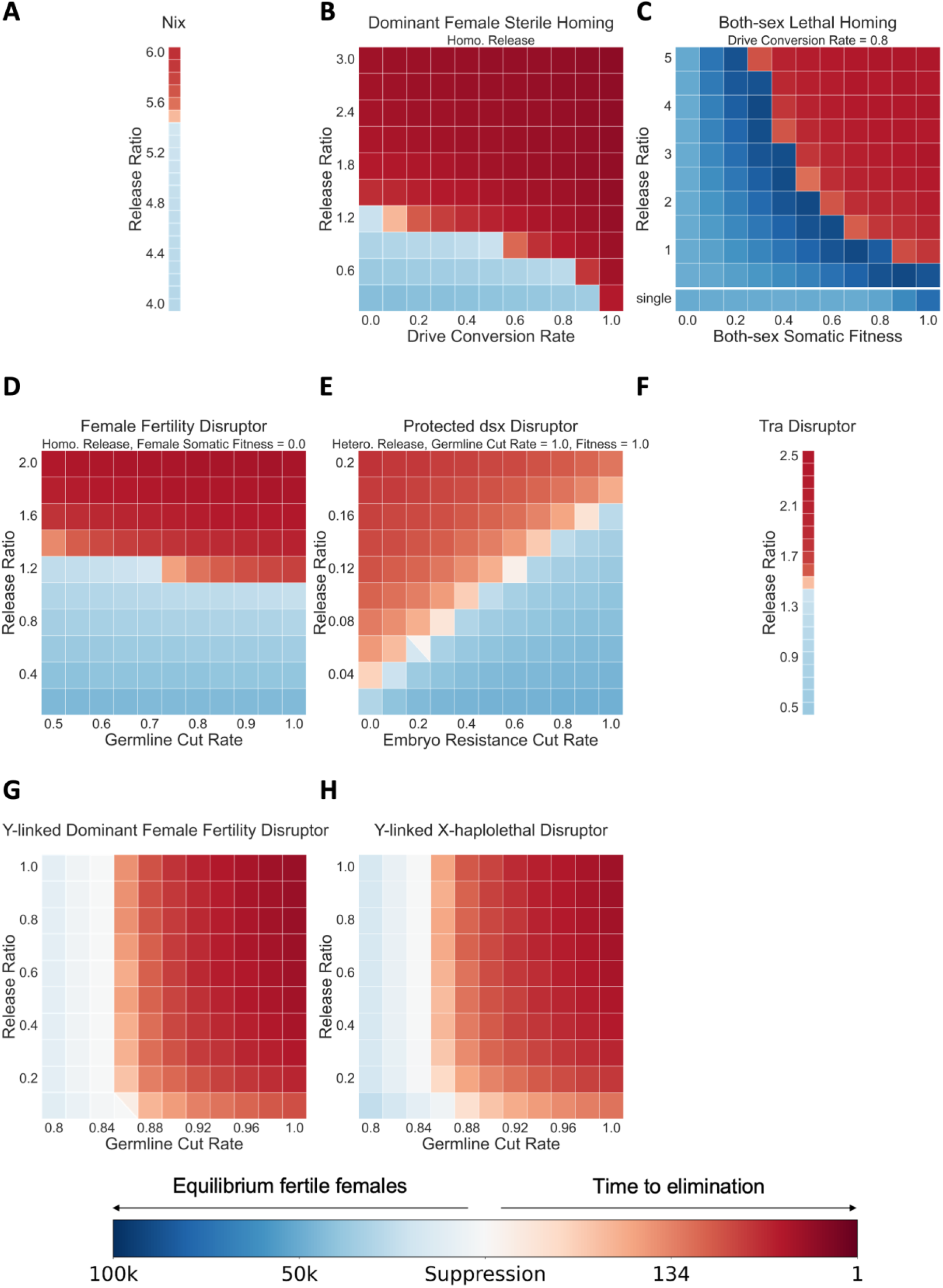
Performances of different systems. Transgenic males with varying suppression systems were released each week at the per-generation release ratios shown into a wild-type population with a carrying capacity of 100,000. With default performance parameters except where indicated, we recorded the time to population elimination or the average number of fertile females after the system reached equilibrium. Squares with two colors indicate that both success and failure happened in simulations. Each point had 10 replicates.

**Figure S3.**
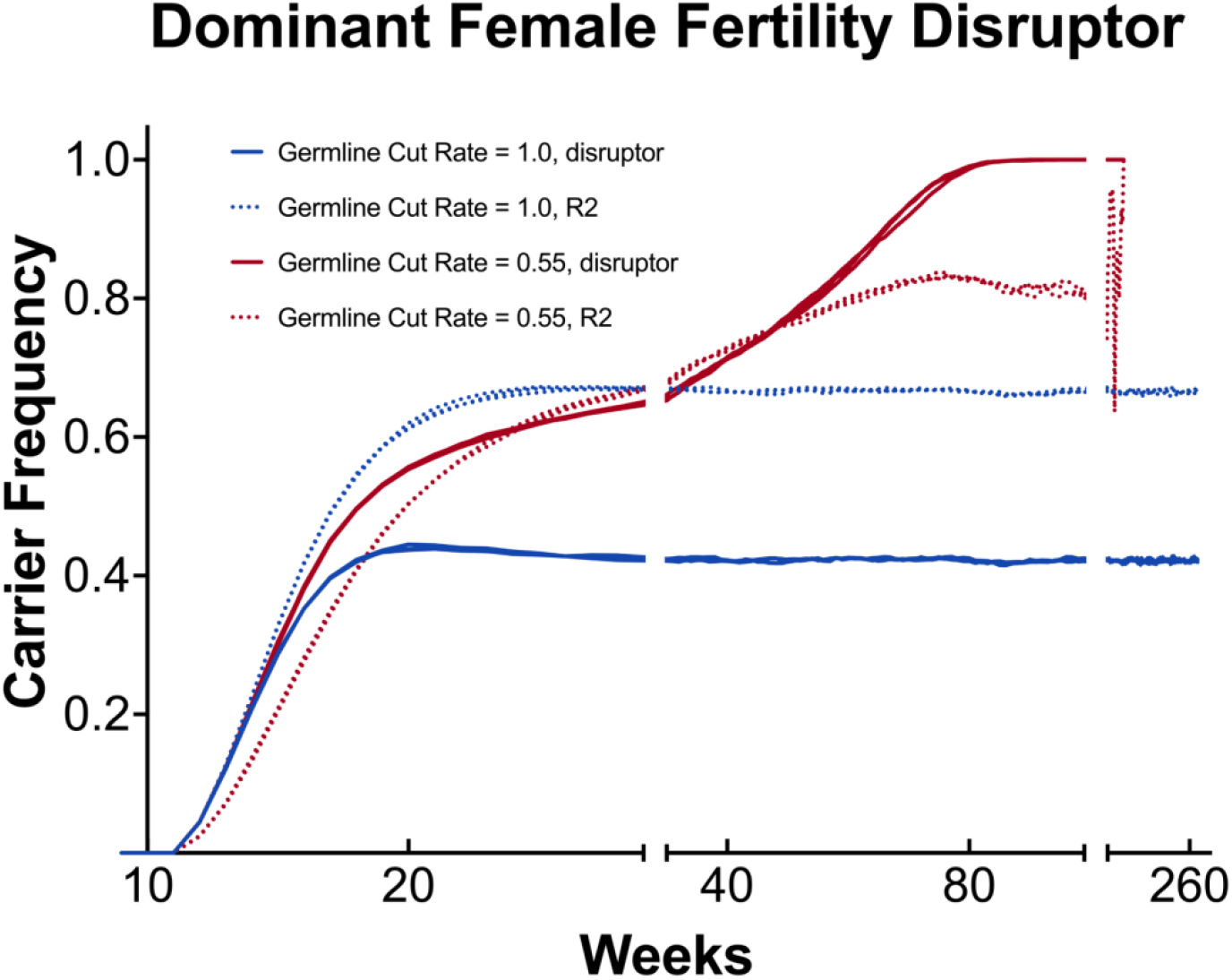
Population dynamics of the dominant female fertility disruptor. Transgenic males with dominant female fertility disruptors were released each week at the per-generation release ratio of 0.5 into a wild-type population with a carrying capacity of 100,000. We recorded the carrier frequency of disruptor and nonfunctional resistance alleles with default parameters and a germline cut rate of 1.0 and 0.55. Three replicates are shown for each cut rate.

**Figure S4.**
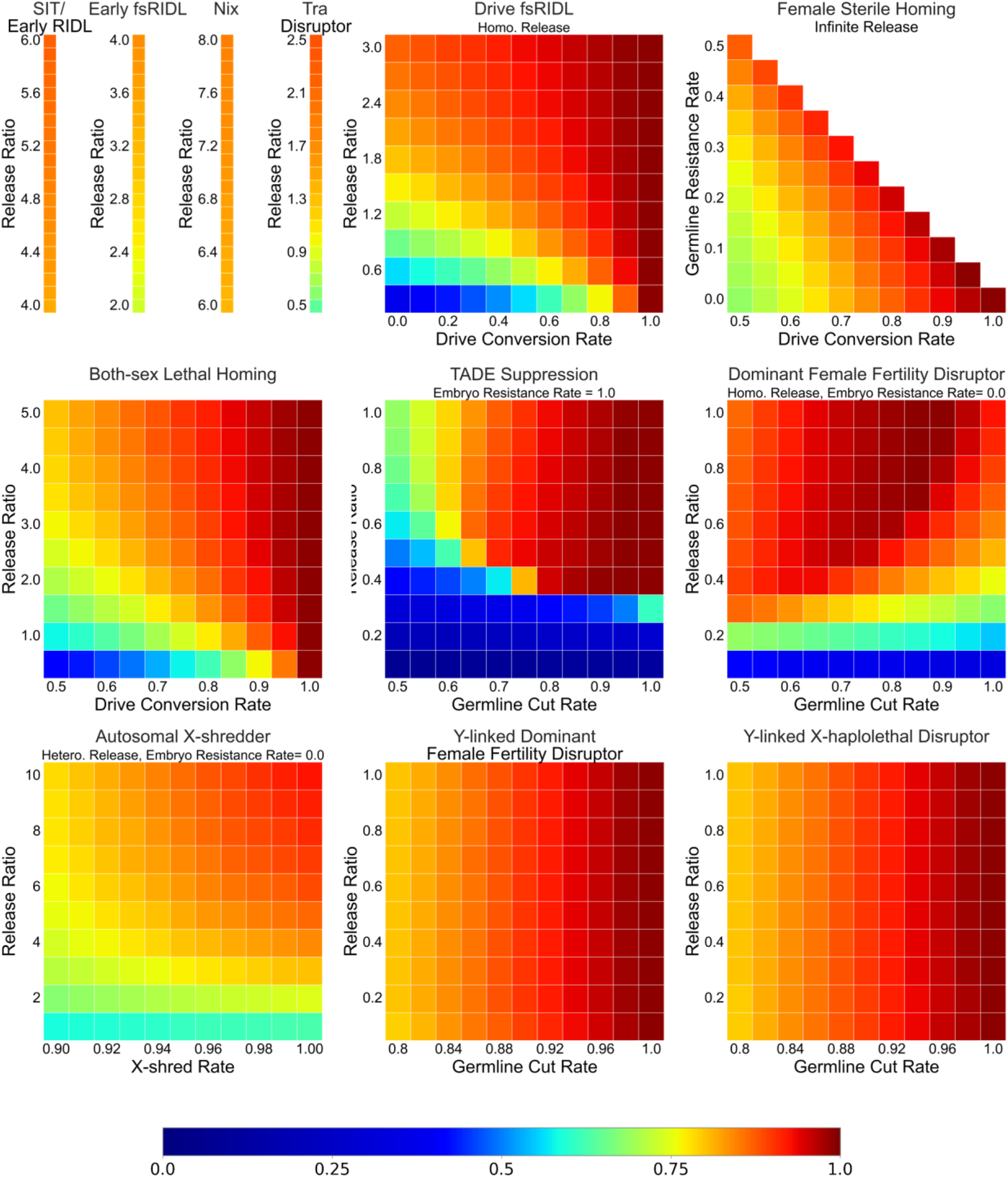
Constant-population genetic load. Transgenic males were released each week at the per-generation release ratios shown into a wild-type population, with the population being kept around its carrying capacity of 100,000. With default performance parameters except where indicated, we recorded the average constant-population genetic load for 50 weeks after the system reached equilibrium. Each point had 5 replicates.

**Figure S5.**
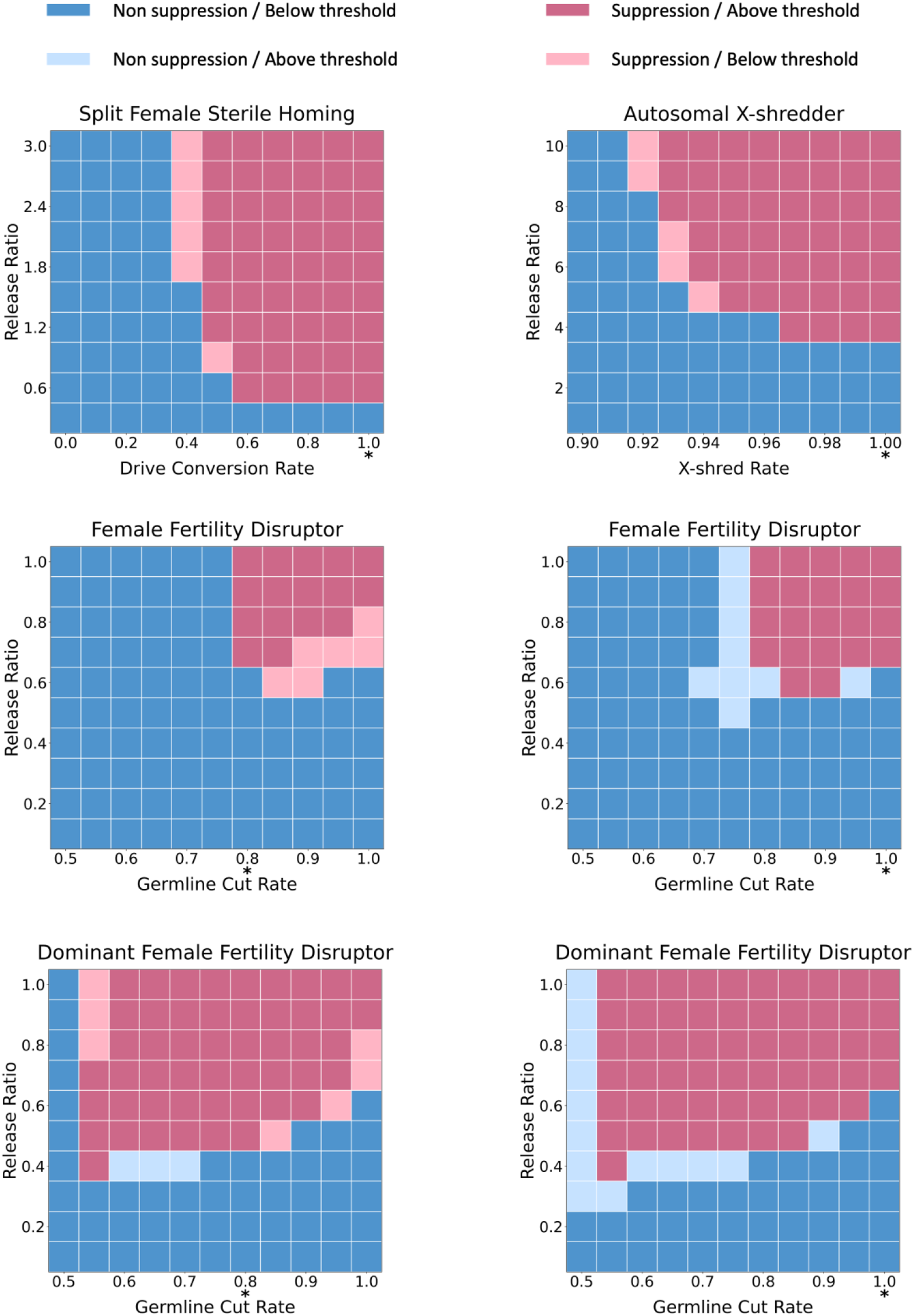
Accuracy of population elimination prediction from constant-population genetic load. Blue and red squares indicate successful and unsuccessful elimination, respectively. Darker colors represent accurate prediction according to constant-population genetic load based on threshold calculated with performance indicated by “*”. Lighter colors represent inaccurate predictions.

**Figure S6.**
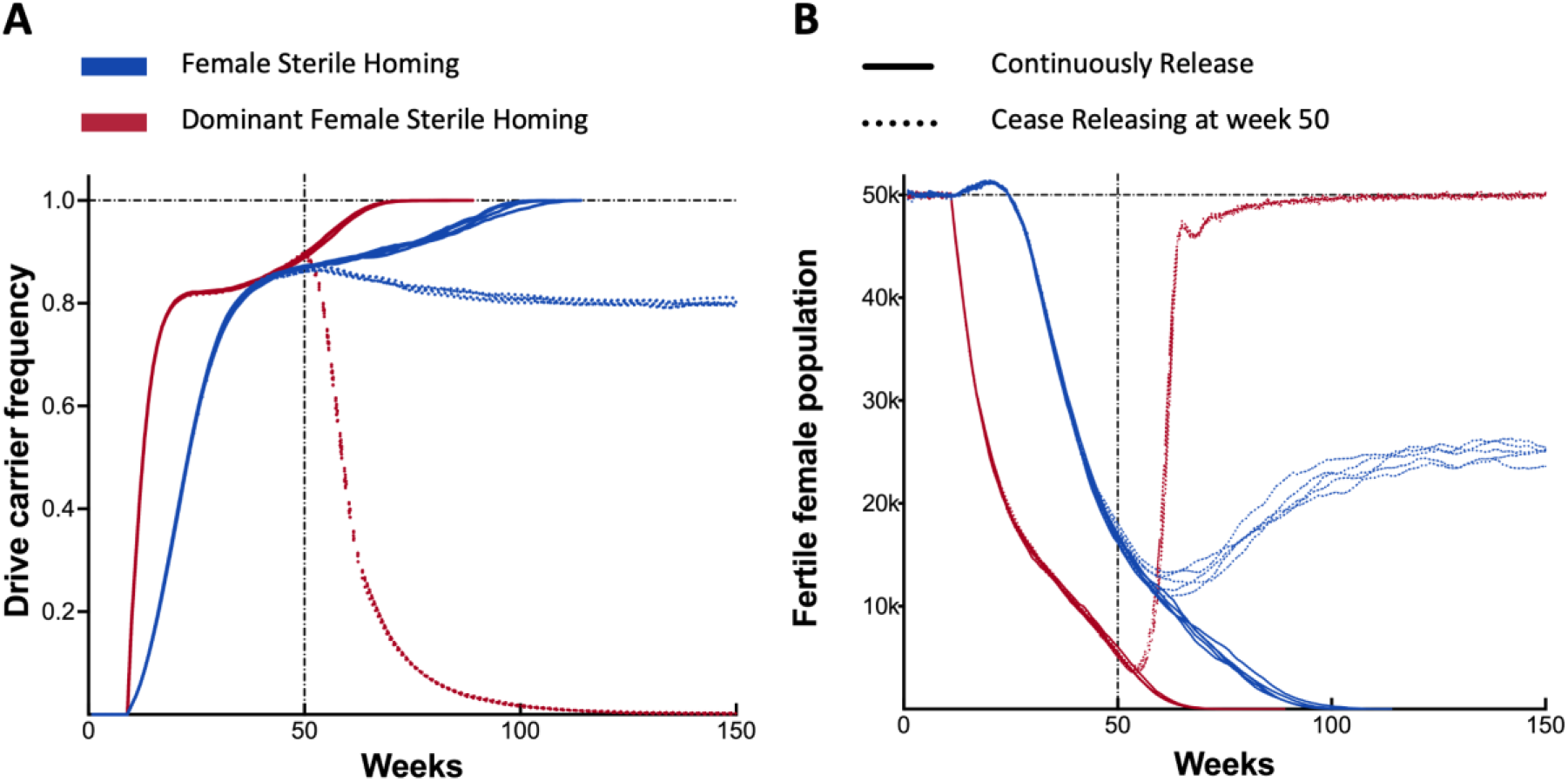
Comparison between self-sustaining and self-limiting suppression systems. Transgenic males with female sterile homing and dominant female sterile homing drives with default parameters were released into a wild-type population with a carrying capacity of 100,000 each week at per-generation release ratios of 0.1 and 1.2, respectively. We recorded the (A) drive carrier frequency and (B) fertile female population when we stopped releases at week 50 and when releases were continued until the end of the simulation. Each scenario shows results from 5 replicates.

